# Nitration of a ribosomal pentapeptide generates a noncanonical precursor for nonribosomal peptide synthesis

**DOI:** 10.1101/2024.07.12.603347

**Authors:** Leo Padva, Lukas Zimmer, Jemma Gullick, Yongwei Zhao, Vishnu Mini Sasi, Ralf B. Schittenhelm, Colin J. Jackson, Max Cryle, Max Crüsemann

## Abstract

Peptide natural products possess a fascinating array of complex structures and diverse functions. Central to this is a repertoire of modified amino acid building blocks, which stem from fundamentally different biosynthesis pathways for peptides of nonribosomal and ribosomal origins. Given these origins, integration of nonribosomal and ribosomal pathways have previously been thought unlikely. Now, we demonstrate that ribosomal biosynthesis generates a key noncanonical 3-nitrotyrosine building block for the nonribosomal synthesis of rufomycin. In this pathway, a biarylitide-type ribosomal peptide is nitrated by a modified cytochrome P450 crosslinking enzyme, with the nitrated residue liberated by the actions of a dedicated protease found within the rufomycin gene cluster before being incorporated into rufomycin by the rufomycin nonribosomal peptide synthetase. This resolves the enigmatic origins of 3-nitrotyrosine within rufomycin biosynthesis and demonstrates unexpected integration of ribosomal peptide synthesis as a mechanism for the generation of noncanonical building blocks within nonribosomal synthesis pathways.

---

Natural product diversity underpins a plethora of biological activities in nature. Within these molecules, peptides form a major class and display diverse functions. Along with their highly variable structures, different activities make peptides important molecules for application in human medicine, for example their use as antibiotics (*1*). Such peptides are produced by two main pathways centered on the use (or exclusion) of the ribosome. Whilst the biosynthetic machinery that produces non-ribosomal peptides – non-ribosomal peptide synthetases (NRPSs) (*2*) – is not limited to the selection of proteinogenic amino acids, ribosomally synthesized and post translationally modified peptide (RiPP) biosynthetic pathways (*3*) require the incorporation of proteinogenic residues that could be seen as a limiting factor for their broader structural diversity. However, recent studies reporting highly modified peptides of ribosomal origin have demonstrated that there is great capacity for diversity within such peptide sequences (*4-9*), thus revealing the biosynthetic scope of these two biosynthetic pathways is closer than was previously envisioned.

Within peptide natural product biosynthesis, one important modification that has been thought to be restricted to non-ribosomal pathways is the nitration of amino acids (*10*). Nitration is a common modification in higher eukaryotes, where it is often a marker of oxidative stress caused by the diffusion of nitric oxide and reactive oxygen species (*11*). The role of nitration in plants is also an area of active investigation, especially given the importance of nitrogen metabolism in these systems and the interplay of soil bacteria and plants in the soil microenvironment. Within peptide biosynthesis pathways, nitration of amino acids has been reported for Trp and Tyr containing peptides, although the direct nitration of amino acids has been restricted to the thaxtomin (tRNA dependent cyclodipeptide synthase) (*12,13*) and rufomycin/ilamycin (NRPS) biosynthesis pathways. (*14,15*) These pathways both utilize a nitric oxide (NO) synthase that generates NO from Arg paired with a cytochrome P450 enzyme that performs nitration (*10*). Cytochrome P450s are a superfamily of diverse oxidative enzymes involved in a range of complex chemical transformations (*16*), and whilst nitration has been reported for P450s, this remains an unusual transformation that lies outside of the classic oxidative chemistry catalyzed by the highly reactive P450 intermediate compound I. (*17*) Such alternate reactivity makes the study of these nitrating P450s of great interest, and whilst the enzyme from thaxtomin biosynthesis (TxtE) has been characterized (*12,13*), the function of the RufO enzyme from rufomycin biosynthesis remains enigmatic.

The rufomycins, also known as ilamycins, are a family of more than 50 characterized congeners of cyclic heptapeptides known for their potent bioactivity against *Mycobacteria*, the causative agent of tuberculosis (*18*). Rufomycins exhibit improved bioactivity compared to existing anti-tuberculosis agents, positioning them as compelling candidates for drug development efforts by targeting the protease ClpC1 in *Mycobacterium tuberculosis* (*19,20*). Structure-activity relationships of the rufomycins have revealed residues central to their antibacterial activity, the most prominent of which is the NO_2_ group of the unique 3-NO_2_-Tyr residue (*21*). The rufomycins are synthesized by a heptamodular NRPS RufT, with the non-proteinogenic L-2-amino-4-hexenoic acid (AHA) moiety synthesized by the polyketide synthase RufE and incorporated by the last module of the rufomycin NRPS (*14,15*) (**Figure 1**). Due to the similarity of the genes *rufNO* to the *txtDE* system, which has been demonstrated to perform nitration of tryptophan in thaxtomin biosynthesis (*13*), the rufomycin 3-NO_2_-Tyr residue has been postulated as being synthesized by an analogous mechanism involving direct nitration of free Tyr (*15*). However, this hypothesis is not supported by reported data, and recent studies suggest RufO instead acts on Tyr bound to a carrier protein domain following selection and activation by the NRPS machinery (*22-24*). The structural characterization of RufO, combined with our interest in biarylitide biosynthesis, led us to postulate an alternate pathway based upon these reports, one involving the unprecedented involvement of a ribosomal biosynthesis pathway to generate the non-proteinogenic 3-NO_2_-Tyr residue for NRPS biosynthesis (**Figure 1**).

**Figure 1.**
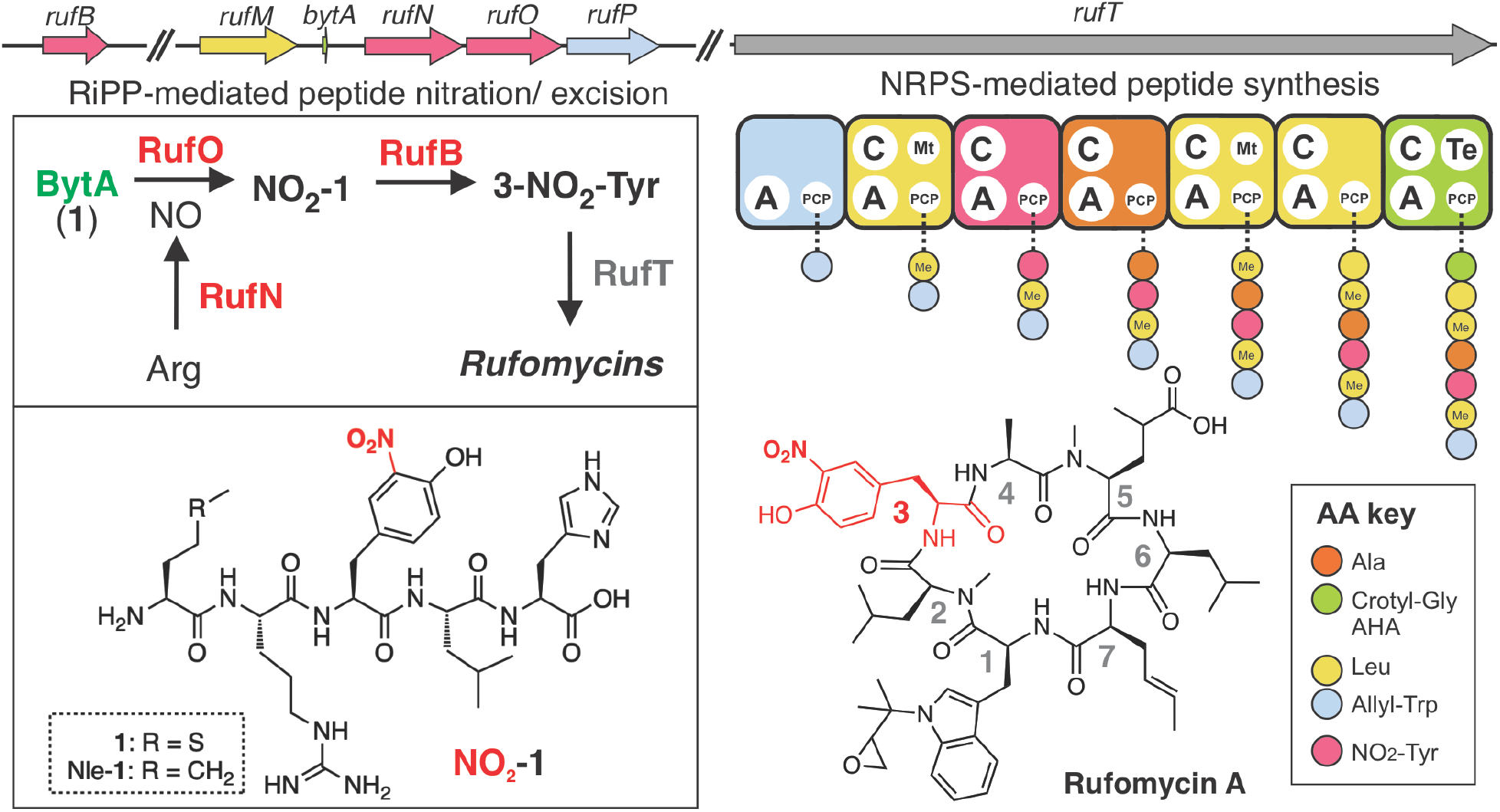
Rufomycin biosynthetic gene cluster (BGC) harbors a RiPP pathway to afford 3-nitrotyrosine. The rufomycin BGC, an NRPS-PKS hybrid, features a RiPP biosynthetic pathway, with *bytA* encoding the pentapeptide MRYLH **1** located upstream of the *rufNO* operon. RufN generates nitric oxide (NO) from Arg, with RufO using NO to nitrate Tyr in **1**. RufB cleaves NO_2_-**1** to release 3-NO_2_-Tyr, which is incorporated into the nonribosomal peptide chain by the RufT A_3_ domain (MLP RufH not shown). Color code links genes to residues in rufomycin; genes to scale except for *rufT;* for complete cluster details see SI; peptide structures shown in **Figure S3**.

The biarylitides are a family of biaryl-linked cyclic tripeptides recently discovered from the actinomycete genus *Planomonospora* (*25*). Amongst the reported expansion of P450s involved in the biosynthesis of crosslinked RiPPs, the biarylitides remain unique due to their biosynthesis being centered on translation of the precursor pentapeptide MRYYH encoded by *bytA*, the smallest known functional gene in the tree of life (*25*). This linear peptide undergoes intramolecular aromatic crosslinking between Tyr-3 and His-5, a transformation catalyzed by the P450 BytO, consistent with other P450 enzymes catalyzing the formation of a range of crosslinked tripeptides (*26-28*). Given our interest in these systems, we were intrigued to find a P450 with 97.97% sequence identity to RufO in one of our in-house strains, *Streptomyces atratus* S3_m208_1 (isolated from a Belgian soil sample (*29*)), located next to a *bytA* homolog encoding a MRYLH pentapeptide upstream of the *rufNO* operon. Remarkably, this biarylitide BGC is found in the center of the rufomycin BGC as well as in all other rufomycin BGCs reported to date (**Figures S1-2**). Given this, we undertook to explore the role of this ribosomal pathway in rufomycin biosynthesis.

## Cytochrome P450 RufO nitrates a RiPP precursor peptide

Our investigations into the function of RufO stemmed from comparisons between its structure (*23,24*) and that of the biarylitide crosslinking P450_Blt_ that we had recently characterized (*26*). Analysis of these structures revealed that they were highly similar (41.6% amino acid ID, C-α RMSD 1.8 Å to 8SPC; **Figure S6**), apart from mutations at a key section of the RufO I-helix involved in oxygen activation (*27*). Indeed, the mutations observed (P450_Blt_: Ser239Val240 to RufO: Val240Pro241; **Figures S7-8**) were indicative of a major alteration in the transformation performed by this P450, compared to what has been seen in the chemistry of other P450s (*30,31*). Given the unexpected presence of a RiPP precursor gene within a NRPS BGC, we next characterized the interaction of RufO with this putative pentapeptide substrate (Nle-RYLH (Nle-**1**); where norleucine (Nle) was used to avoid Met oxidation; **Figures 2, S9-10**). Binding experiments showed that the affinity of Nle-**1** for RufO was comparable to that of P450_Blt_ (0.84 µM vs 2.1 µM; **Figure S10**), with a shift in the soret maximum on binding to 410 nm. Such tight binding was somewhat unexpected given the bulky residue Val replaces the active site Ser in P450_Blt_ that is involved in key H-bonding interactions with the substrate peptide in P450_Blt_ (*26*). The effects of this alteration in binding were investigated by further analyzing the interactions of **1** with P450_Blt_ and the S239V/V240P mutant using molecular dynamics simulations, which showed **1** adopting a conformation in the mutant in which the C-terminal residues of Nle-**1** orient away from the I-helix. Little alteration in position was observed with the Tyr-3 residue despite the mutation of Ser239 to valine (**Figure S11**).

**Figure 2.**
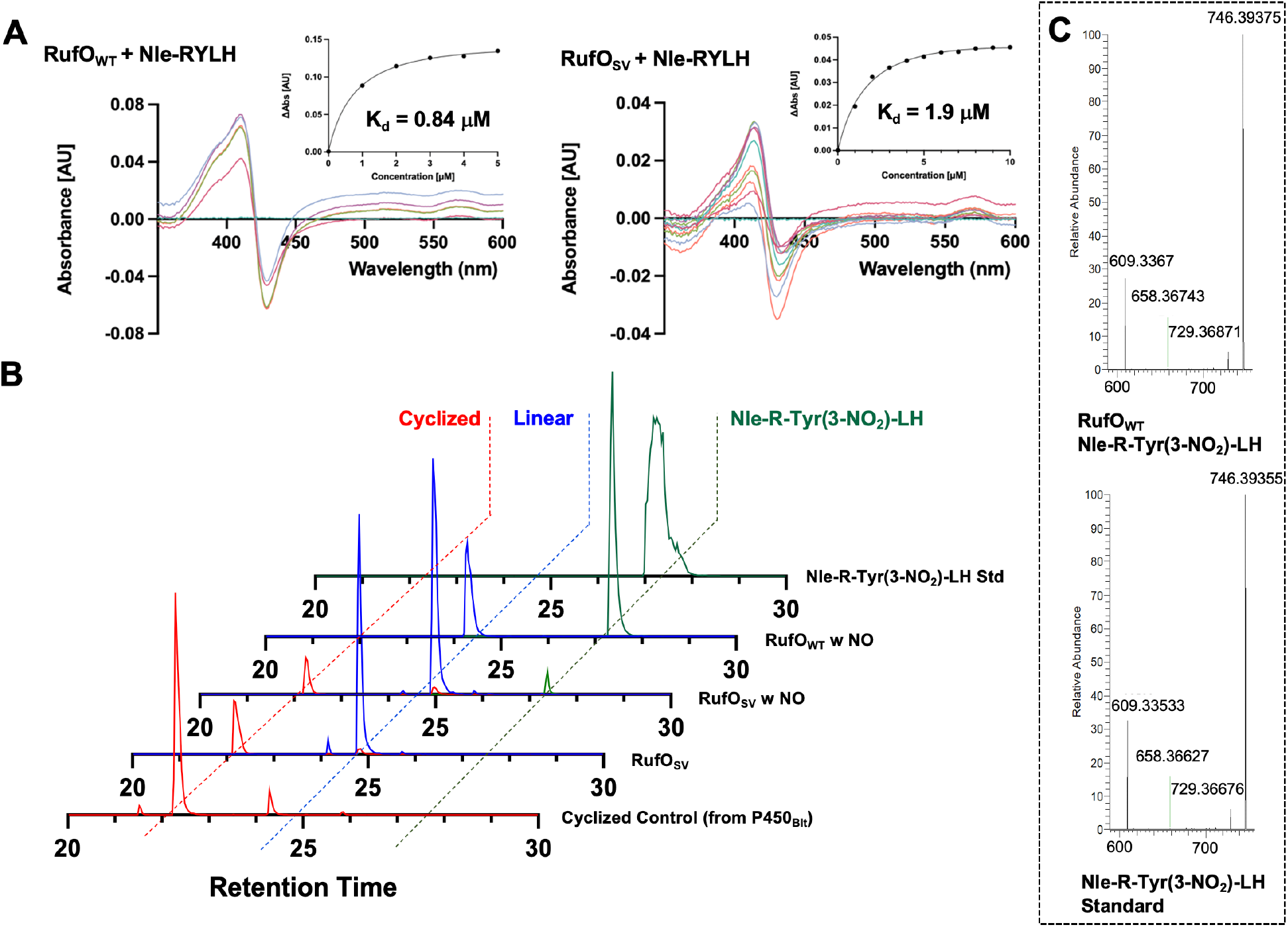
*In vitro* characterisation of RufO-catalysed nitration of Nle-1. Substrate binding traces with affinities for RufO and RufO_SV mutant for Nle-**1** (A); turnover traces for RufO, RufO_SV and P450_Blt_ with Nle-**1** (B); and comparison of MS^2^ fragmentation of nitrated Nle-**1** from RufO turnover with synthetic standard Nle/NO_2_-**1** (C).

Next, we turned to P450-catalysed turnover experiments to clarify the activity of RufO (**Figures S12-20**). Attempts to crosslink Nle-**1** using traditional redox partner proteins with RufO were not successful, which stands in contrast to the high levels of crosslinking seen with P450_Blt_ (*28*). Given this, we explored the potential for RufO to nitrate peptide Nle-**1**. Using protocols reported for the diketopiperazine nitrating P450 TxtE (with nitrating reagent DEANO (*13)*) we saw high levels of nitration, with HR-MS_2_ and comparison to authentic standards demonstrating nitration as occurring exclusively on the Tyr-3 residue of Nle-**1** (**Figure 2**). We were also able to affect nitration using dithionite in place of redox partner proteins (*28*), which supports the postulated mechanism of this reaction as occurring prior to the formation of the typical P450 reactive intermediate (Compound 1) (*28,31*). Substitution of His for Trp in a MRYLW substrate peptide demonstrated crosslinking activity from RufO without nitration, supporting the crucial nature of the altered peptide binding mode seen with **1** and the P450_Blt_ mutant in MD simulations regarding the peptide C-terminus. Having seen the switch in chemistry occurring with RufO, we tested the effect of mutation of the altered I-helix residues in RufO to those seen in the crosslinking P450_Blt_. Whilst a single Val240 to Ser mutant was unable to be expressed in soluble form (suggesting the mutation was structurally destabilizing), Val240Ser/Pro241Val (RufO_SV) could be expressed, albeit with somewhat reduced yield. The effect of this double mutation was reversion of activity from nitration of Nle-**1** to crosslinking (20% conversion; 1:5 when DEANO was present), although curiously the rate enhancement seen with the addition of DEANO in wildtype RufO was maintained in this mutant despite the lack of nitration. Low levels of cyclized and nitrated **1** was also detected in these assays, showing an ability for the enzyme to perform sequential transformations. The RufO mutations (Ser239Val/Val240Pro) were also incorporated into P450_Blt_ to make it resemble RufO, although this mutant was unable to catalyze nitration or crosslinking. These results demonstrate the importance of I-helix residues in controlling P450 catalysis in biarylitide pathways and implied an unexpected route to generate 3-NO_2_-Tyr in rufomycin biosynthesis.

## RufT module 3 A-domain accepts 3-NO_2_-Tyr

With evidence supporting the formation of 3-NO_2_-Tyr via RufO-mediated activity towards the peptide Nle-**1**, we next sought to validate the incorporation of 3-NO_2_-Tyr by the A_3_ adenylation domain of the NRPS enzyme RufT, since this activity has not been previously determined. Structural modelling using Alphafold (*32*) and analysis of the substrate binding pocket together with comparison to Tyr-accepting A-domains pointed to an enlarged substrate binding pocket due to a Cys to Gly mutation in the penultimate position of the A-domain specificity code, providing additional space for the pendant NO_2_ group and allowing favorable interactions with an unusual Lys residue found in the A_3_ domain and made accessible by this mutation (**Figure 3A, S21-22**). We designed constructs of A_3_ with varying lengths and including regions of the adjacent condensation (C) or peptidyl carrier protein (PCP) domains, which were co-expressed with the MbtH-like protein (MLP) RufH encoded in the rufomycin BGC (*33*). One A-PCP construct was successfully co-expressed with RufH and was purified to evaluate activation activity using an γ^18^O_4_-ATP exchange assay (*34*). Somewhat surprisingly, both L-Tyr and 3-NO_2_-Tyr were adenylated at similar rates (**Figure 3B**), indicating promiscuity of this A-domain; this also agrees with the large number of natural and mutasynthesis-derived rufomycin analogs with variations in the Tyr position (*35*). Such promiscuity in Tyr-activating domains has previously been reported for the A_6_ domain from teicoplanin biosynthesis (*36*) and suggests possible analogous proofreading by the RufT C_2_ domain during peptide extension. These results supported the direct activation of 3-NO_2_-Tyr by the RufT A_3_ domain and so we turned to *in vivo* assays to clarify the biosynthesis of 3-NO_2_-Tyr.

**Figure 3.**
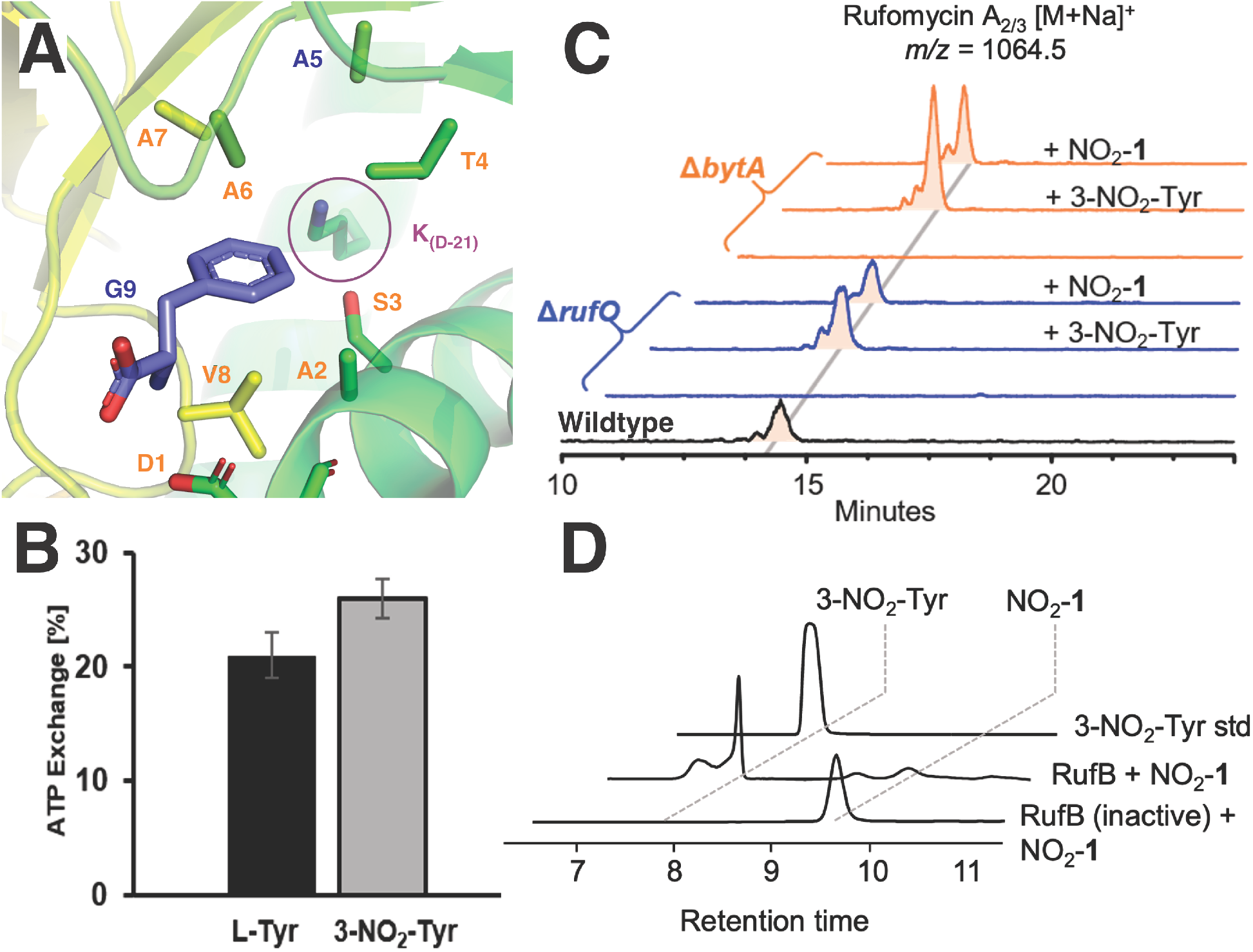
Evaluating incorporation of NO_2_-Tyr into rufomycin and release from NO_2_-1. Alphafold model of the RufT A_3_ domain with overlay of Phe from PheA A-domain (PDB: 1AMU) showing key selection residues 1-9 plus unusual Lys residue (A); γ^18^O_4_ -ATP exchange assay of substrate activation for L-Tyr and 3-NO_2_-Tyr by RufT A_3_ domain (B); Rufomycin A2/3 production in *S. atratus* S3_m208_1 wildtype (black) Δ*rufO (blue*) and in Δ*bytA (orange*) mutant strains, with rufomycin biosynthesis restored in the mutants on supplementation with either 3-NO_2_-Tyr or peptide NO_2_-**1** (C); RufB specifically digests NO_2_-**1** to release the central, modified amino acid 3-NO_2_-Tyr (D). Retention times shown in minutes.

## Role of the BytA peptide in rufomycin biosynthesis

We used *S. atratus* S3_m208_1 to test its ability to produce rufomycins under laboratory conditions and further to examine the role of *bytA*, which we reasoned must be fundamental for generation of 3-NO_2_-Tyr and thus for rufomycin biosynthesis. We designed a knockout plasmid for homologous recombination, incorporating a 12 bp deletion in *bytA* and cloned this into pYH7 (*37*). After conjugation to *S. atratus* S3_m208_1 and selective passaging, we obtained clones showing desired recombination. Crucially, *S. atratus* S3_m208_1 Δ*bytA* lost the ability to produce the rufomycins, supporting the essential nature of *bytA* for rufomycin biosynthesis. Feeding 3-NO_2_-Tyr or NO_2_-**1** to cultures of the mutant could complement this deletion mutant and restore rufomycin biosynthesis, confirming the essential role of the MRYLH peptide **1** (**Figure 3C, S23**). We then used CRISPR-Cas9-cBest (*38*) to introduce a specific stop codon into *rufO* and generated the *S. atratus* S3_m208_1 Δ*rufO* mutant. This mutant also showed a loss of rufomycin biosynthesis in the same manner as its substrate peptide **1** encoded by *bytA*. Supplementation of 3-NO_2_-Tyr or NO_2_-**1** to the Δ*rufO* mutant could re-establish rufomycin production, further supporting the role of this biarylitide RiPP pathway in rufomycin biosynthesis (**Figure 3C, S24**).

## RufB is an aminopeptidase that cleaves the BytA pentapeptide 1

The availability of 3-NO_2_-Tyr as a substrate for the biosynthesis of rufomycin requires the release of the modified amino acid from the precursor pentapeptide NO_2_-**1**. Leader peptide removal during RiPP biosynthesis is a diverse process and can be catalyzed by either specific or non-specific proteases within the cytoplasm, during secretion or extracellularly (*39*). Given the essential role of the *bytA* encoded peptide for rufomycin biosynthesis, we hypothesized that a specific mechanism would be involved in the release of 3-NO_2_-Tyr from the pentapeptide NO_2_-**1**. Inspection of the rufomycin gene cluster revealed the presence of the RufB enzyme annotated as an α/β fold hydrolase containing a serine aminopeptidase domain (IPR022742). A reported deletion mutant of the *rufB* homolog *ilaC* showed production of ilamycin (rufomycin) was halved compared to the wildtype producer (*14*), although the exact function of this enzyme was not determined. To test our hypothesis that RufB was involved in 3-NO_2_-Tyr release from NO_2_-**1**, we expressed RufB fused with an N-terminal maltose binding protein tag in *E. coli* (*40*) that was purified and used in digestion assays with NO_2_-**1**. HPLC-MS analysis showed that RufB completely digested NO_2_-**1** for concomitant release of 3-NO_2_-Tyr (**Figure 3D, S25**). These results reveal the role of RufB as a BGC-encoded peptidase acting upon the nitrated peptide NO_2_-**1**, a requirement for the release of 3-NO_2_-Tyr from the BytA peptide and allowing subsequent incorporation by the rufomycin NRPS machinery.

## Discussion

Natural product biosynthesis demonstrates remarkable examples of biosynthetic ingenuity. Whilst specific biosynthetic pathways can already produce compounds with great chemical complexity, synergising assembly lines can afford yet even more diversity in such compounds. While increasing examples of such synergy have been identified across pathways, one example that has proved more elusive is the integration of peptide biosynthesis from both ribosomal and non-ribosomal pathways. Here, we have investigated the generation of 3-NO_2_-Tyr in rufomycin biosynthesis and could demonstrate how these pathways can be synergized to deliver non-canonical amino acid building blocks in peptide biosynthesis. The rufomycin pathway reveals that P450 enzymes within RiPP pathways can perform specific nitration on peptide substrates with only minimal changes to the active site. The evolution of a peptide crosslinking P450 – of which 20 classes have recently been identified (*41*) – into one capable of selective nitration shows parallels to the ability of radical SAM dependent enzymes to rapidly diversify their function with minimal changes to the enzyme (*42*). The identification of mutations to key P450 residues as a fingerprint for alternate catalysis (*31*) offers important clues to identify examples of diversified P450 function, whilst demonstrating the power of the P450 active cycle to access different intermediates during catalysis and underlining the versatility of P450s in biosynthetic pathways.

Finally, the lack of detection of the *bytA* gene in the rufomycin BGC demonstrates the challenges facing biosynthetic discovery due to the small size of such genes and the current lack of bioinformation detection pipelines for these minimal RiPPs. Recent examples of combining ribosomal biosynthesis with additional pathways include the peptide-amino acyl tRNA ligase (PEARL) enzymes, which act on ribosomally synthesized peptides to catalyze the attachment of amino acids on the C-terminus of a scaffold peptide in an ATP- and tRNA-dependent manner and that are capable of previously unprecedented biosynthetic transformations (*43-45*). The biosynthesis of rufomycin represents a new instance of comparable biosynthetic ingenuity and is the first report of a complete RiPP biosynthesis pathway merged within nonribosomal peptide biosynthesis. The fact that this RiPP pathway serves to synthesize a rare amino acid building block for an NRPS – a system classically prized for its ability to install nonproteinogenic amino acids – and that the P450 enzyme central to the pathway has evolved to perform a new catalytic function to accomplish this – once more highlights how nature has developed novel solutions for complex biosynthetic problems. It also demonstrates the importance of continued study to identify and characterize such novel examples of biocatalysis, of which there are no doubt many more potent examples to discover.

## Acknowledgments

We thank Aurélien Carlier (INRAE, Toulouse) for providing *S. atratus S3_m208_1*; Tilmann Weber (DTU Copenhagen) for providing CRISPR-Cas9-cBest plasmids; Mathias Hansen (Monash) & James De Voss (UQ) for helpful discussions; Stefan Kehraus (Uni Bonn) for LCMS measurements.

## Author contributions

LP: *in vitro* P450 assays, *in vivo* experiments, *in vitro* RufB and A domain assays, figure & manuscript preparation. LZ: gene cluster analysis, *in vivo* experiments, *in vitro* RufB and A domain assays. JG, YZ: synthesis, *in vitro* P450 assays, analysis. RS: HRMS analysis. VMS, CJJ: MD simulations. MJC & MC: analysis, figure & manuscript preparation.

## Data and materials availability

See Supplementary Information. The mass spectrometry data for *in vitro* P450 enzyme assays have been deposited to the ProteomeXchange Consortium via the PRIDE partner repository with the dataset identifier PXD053820.

