## Supplementary Information for "Nitration of a ribosomal pentapeptide generates a noncanonical precursor for nonribosomal peptide synthesis" (from another version of this preprint: version 1, 2024-07-13)

| | |
|---|---|
| Materials and Methods | Page 2 |
| Supplementary Tables | Page 8 |
| Supplementary Figures | Page 13 |
| Supplementary References | Page 37 |

### Materials and Methods

#### Materials

Fmoc-protected amino acids (Sigma-Aldrich), 2-chlorotrityl chloride resin (GL Biochem), dichloromethane (Thermo Fisher), *N,N*-dimethylformamide (Ajax Finechem, Thermo Fisher), methanol (Scharlau), trifluoroacetic acid (Oakwood chemical), triisopropyl silane (Sigma-Aldrich), diethyl ether (Sigma-Aldrich), OxymaPure (Novabiochem), N,N'-diisopropylcarbodiimide (Sigma-Aldrich), $NaH_2PO_4$ (Sigma-Aldrich), $Na_2HPO_4$ (Sigma-Aldrich), formic acid (FA, Sigma-Aldrich), 4-(2-hydroxyethyl)-1-piperazineethanesulfonic acid (Sigma-Aldrich), isopropylthio-β-galactoside (Promega), tryptone (MP Biochemicals), yeast extract (MP Biochemicals), glycerol (MP Biomedicals). imidazole (Sigma-Aldrich), Tris-HCl (Sigma-Aldrich), δ-aminolevulinic acid (Sigma-Aldrich), NaCl (Sigma-Aldrich), glucose (Sigma-Aldrich), glucose dehydrogenase (Sigma-Aldrich), nicotinamide adenine dinucleotide (Sigma-Aldrich), diethylamine NONOate (Sigma-Aldrich), kanamycin sulfate (Sigma-Aldrich), gentamycin sulfate (Sigma-Aldrich), *E. coli* 5$\alpha$ competent cells (NEB), *E. coli* BL21 (DE3) competent cells (NEB), *E. coli* 10β competent cells (NEB), *E. coli* ArcticExpress (DE3) competent cells (Agilent), Ni-NTA agarose (Macherey-Nagel), Precision Plus Protein™ Dual Colour Standards (Bio-Rad), Amicon® Ultra-0.5 mL centrifugal filters units (30 kDa MWCO; Sigma-Aldrich)), SPE Bond elute (Agilent), Q5-Polymerase (NEB), Phusion polymerase (NEB), *Taq* ligase (NEB), 3-Nitro-L-Tyrosine (Sigma-Aldrich).

#### Methods

##### Identification and comparison of ilamycin/rufomycin BGCs

Amino acid sequences of RufN (BBA20961.1) and RufO (BBA20962.1) encoded in the rufomycin BGC from *Streptomyces atratus* ATCC 14046 were used as queries to detect BGCs harboring these two genes (*46*). For the analysis of homologous BGCs, cblaster was used with the default settings (*47*). The genome sequences of the obtained hits were collected from the NCBI database and were analysed using antiSMASH 7.0 to determine the location of *rufN*/*rufO* and whether the respective organisms possess a rufomycin/ilamycin BGC (*48*). The region GenBank files of all gene clusters harbouring *rufN*/*rufO* were then collected for alignment and visualization of the gene clusters using CAGECAT (*49*). This analysis was performed using a minimum alignment sequence identity of 0.45.

##### Synthesis of *L*-tyrosine-(phenyl-3,5-$d_2$) and Fmoc-L-tyrosine-(phenyl-3,5-$d_2$)

The syntheses of L-tyrosine-(phenyl-3,5-$d_2$) and Fmoc-L-tyrosine-(phenyl-3,5-$d_2$) were performed according to the previously reported protocol (*50*). All spectra matched that previously reported (*50*).

##### Peptide synthesis

Peptide synthesis was performed manually at 0.1 mmol scale using DIC and Oxyma Pure as coupling reagents. 2-chlorotrityl chloride resin (97 mg) was washed with DCM (3 x 2 ml), swelled in DCM (2 ml, 30 min), and washed again with DCM (3 x 2 ml). The first Fmoc protected amino acid (0.15 mmol) was coupled to the resin overnight using DIEA. The unreacted chlorotrityl groups were capped using MeOH by washing with 3x 2 ml DMF/DIEA/MeOH (7:2:1). After the Fmoc removal step using 20% piperidine in DMF, Fmoc-amino acid coupling (3 eq) was performed using DIC (3eq) and Oxyma Pure (6 eq) at 50 °C for 10 min. The deprotection and coupling reaction was repeated until the sequence was finished, and a final deprotection was performed to remove the Fmoc protecting group of the final amino acid. The cleavage of peptide from resin was performed using 5 mL of TFA/TIS/$H_2O$ (95: 2.5: 2.5, v: v': v'') solution with shaking for 1 h at room temperature (2 h for Arg(Pbf) containing sequence). The separation of resin from the solution was achieved by filtration. After washing the resin with TFA (2 x 1 mL), the filtrate was concentrated under a stream of $N_2$ to ~1 mL, followed by peptide precipitation with pre-cooled diethyl ether (20 mL), centrifugation in a spark-free centrifuge (Spintron, GT-175) and washing with diethyl ether (3x 10 mL). Peptides were analysed by LC-MS (Shimadzu) using a gradient of ACN (0.1% FA) from 0% to 50% over 30 min. For structures see **Supplementary Figure S3**.

**Site-directed mutagenesis**

Point mutations in this study are generated by site-directed mutagenesis. Primer pairs containing the desired mutation are designed and purchased from Sigma-Aldrich. These primer pairs are used in PCR to generate mutated plasmid containing nicks using Phusion® High-Fidelity PCR Master Mix with HF Buffer (NEB) or Q5® Hot Start High-Fidelity DNA Polymerase (NEB). The PCR products were treated either with *Dpn*I endonuclease (NEB) to digest the methylated parental DNA template when sticky ends were generated, or with KLD Enzyme Mix (NEB) perform the ligation as well as template digestion when blunt ends are generated. After the reaction, the mixture was used for transformation of NEB 10-ß competent *E. coli* cells. Following the overnight incubation, plasmid DNA from individual colonies was isolated and sequenced to confirm the correct construction of target proteins.

**Expression and purification of P450 enzymes and mutants**

The expression and purification of $P450_{Blt}$ and mutants were performed as previously described (*51*). The expression and purification of RufO also used the same protocol (*51*).

**Peptide cyclisation and nitration assay**

The cyclisation assay was performed as previously described (*51*). Briefly, HEPES (50 mM), NaCl (50 mM), P450 (0.5 μM), PuR (0.5 μM), PuxB A105V mutant (2.5 μM) (*52*), glucose (0.33%), glucose dehydrogenase (0.033 mg/mL), peptide substrate (25 μM) and NADH (2 mM) were combined and incubated overnight at 30 °C with 300 rpm shaking. For the nitration reaction nitric oxide was also supplied in the form of diethylamine NONOate (DEANO, 0.5 mM). For the reaction with sodium dithionite, no PuR, PuxB A105V, glucose or glucose dehydrogenase were added and sodium dithionite (100 μM) was added instead. Purification was then performed by solid-phase extraction (Bond elute, Agilent) that had been activated with 500 μL of MeOH (0.1 % FA) and equilibrated with 1 mL of water (0.1% FA). The reaction was diluted to 1 mL using water (0.1% FA) before being applied to the column, subsequently washed with 1 mL of water (0.1 % FA) and finally eluted with 600 μL of MeOH (0.1 % FA). The methanol elution fraction was dried using a concentrator (Eppendorf), before being re-dissolved in 50 μL of ACN/$H_2O$ (1:4, 0.1 % FA) before LCMS analysis. All LCMS analyses were carried out on a Shimadzu high-performance liquid chromatography system coupled to mass spectrometer LCMS-2020 (ESI, operating both in positive and negative mode) equipped with an SPD-20A Prominence photo diode array detector and an LC-20AD solvent delivery module. Analytical separations were performed using an Agilent ZORBAX 300SB-$C_{18}$ 5 μm column. All cyclic pentapeptide products were confirmed by HRMS analysis.

**NADH Consumption Assay**

The consumption of NADH by P450s was measured using a Jasco V-750 spectrophotometer at 30 °C. A cyclisation or nitration assay was set up as described above with no addition of glucose, glucose dehydrogenase, NADH and the peptide substrate. A blank measurement was taken prior to the addition of NADH to a final concentration of 100 μM. The absorbance was then measured at 340 nm for two minutes to obtain a background slope ($Abs_{back}$) before peptide substrate (25 μM) was added and the absorbance measured for three minutes. The slope of the curve after NADH addition was then determined ($Abs_{samp}$). The $V_{max}$ and $K_{cat}$ values were calculated using the molar extinction coefficient of NADH at 340 nm and the following equations: $V_{max} = (Abs_{samp} - Abs_{back})/0.00622*1$ and $K_{cat} = V_{max}/[P450]$.

**P450 CO difference spectra**

Spectra were obtained using a Jasco V-750 spectrophotometer at 30°C. The enzyme was diluted to 2.5 μM using Tris-HCl (50 mM, pH 7.4) and a baseline spectrum obtained. CO was generated by the addition of a small volume of disodium boranocarbonate, and the P450 reduced using 20 μL of freshly prepared saturated sodium dithionate solution. The UV-Vis spectra of CO-complex was then measured until the spectra was stable.

**P450/substrate binding determined via UV-Vis spectroscopy**

The interaction of substrates with P450s were measured using UV-visible absorbance spectroscopy on a Jasco V-750 spectrophotometer at 30 °C. The P450 enzyme was diluted to 3 μM using Tris-HCl (50 mM, pH 8.0) and split into two cuvettes. After being equilibrated to 30 °C, a baseline measurement was obtained, and the initial spectra measured between λ = 350 and 600 nm. The substrate was then added into the sample cuvette in a stepwise manner, with the same amount of buffer added into the reference cuvette. The spectra were measured each step after a 2 min equilibration period and normalised against the 600 nm absorbance. The resulting amplitudes of the different spectra were extrapolated ($\Delta A = A_{max} - A_{min}$) and plotted against the ligand concentration. Curve fitting of the resulting data points was performed using GraphPad Prism 9 with a one-site binding model: $Y=B_{max}*X/(K_D+X) + NS*X + Background$ ($Y$ = delta Abs, $B_{max}$ = the maximum specific binding, $K_D$ = equilibrium dissociation constant, $NS$ = slope of nonspecific binding, *Background* = nonspecific binding with no added ligand).

**HRMS analysis of *in vitro* turnovers**

High-resolution mass spectrometry measurements were performed on an Orbitrap Fusion mass spectrometer (Thermo Scientific) coupled to a Dionex UltiMate 3000 RSLCnano system equipped with a Dionex UltiMate 3000 RS autosampler, an Acclaim PepMap RSLC analytical column (75 μm x 50 cm, nanoViper, C18, 2 μm, 100 Å; Thermo Scientific) and an Acclaim PepMap 100 trap column (100 μm x 2 cm, nanoViper, C18, 5 μm, 100 Å; Thermo Scientific). The peptides were separated by increasing concentrations of 80% acetonitrile / 0.1% formic acid at a flow of 250 $nL.min^{-1}$ over 30 min. The instrument was operated in alternating data-dependent acquisition (DDA) and parallel reaction monitoring (PRM) cycles in such that 5 $MS^2$ scans were triggered per survey $MS^1$ scan followed by several targeted $MS^2$ scans to ensure fragmentation of predefined, sample-dependent *m/z* precursors. Each survey $MS^1$ scan (300–1800 *m/z*) was acquired with a resolution of 240,000 and a normalised AGC (automatic gain control) target of 200%. Dynamic exclusion was set to 10 seconds after one occurrence. The 5 most intense ions were selected for HCD fragmentation (fixed collision energy mode, 24% HCD Collision Energy) with a resolution of 15,000, a normalised AGC target of 200% and a fixed first mass of 100 *m/z*. Subsequent targeted ms2 scans were acquired with essentially identical settings. The raw data files were analysed with QualBrowser (XCalibur 3.0.63, Thermo Scientific) to view spectra and to generate extracted ion chromatograms.

**Molecular Dynamics Simulations**

***Protein Preparation.*** All the modelling work, including MDS, was performed in Maestro (Schrödinger Release 2021-3: Maestro, Schrödinger, LLC, New York, NY, 2021). Protein preparation wizard (Schrödinger Release 2021-3: Protein Preparation Wizard; Epik, Schrödinger, LLC, New York, NY, 2021; Impact, Schrödinger, LLC, New York, NY; Prime, Schrödinger, LLC, New York, NY, 2021) was accessed to perform structural edits on one monomer of the protein, which involved correcting missing hydrogens and bond orders, adding missing sidechains and zero-order bonds to heme $Fe^{2+}$ ion and finally assigning protonation states at a pH of 7.4 using Epik. The hydrogen bond network of the prepared structure was further optimised using PROPKA and minimised by converging the het atoms at an RMSD of 0.3 Å.

***Molecular modelling studies.*** The Build module (Schrödinger Release 2021-3: Maestro, Schrödinger, LLC, New York, NY, 2021) was utilised to selectively mutate the Ser239 and Val240 to Val239 and Pro240 in the $P450_{Blt}$ – MRYLH complex, respectively. The Ser239Val/Val240Pro mutant model was energy-minimized using the Prime (Schrödinger Release 2021-3: Prime, Schrödinger, LLC, New York, NY, 2021) module to generate the starting representative models for molecular dynamics simulations.

***Molecular dynamics simulations (MDS).*** The representative model was assigned a simulation box using the System Builder module (Schrödinger Release 2021-3: Desmond Molecular Dynamics System, D. E. Shaw Research, New York, NY, 2021). The TIP4P water model was utilized, with periodic boundary conditions involving an orthorhombic box shape using the buffer method (12 × 12 × 12 Å). After minimizing the volume, the net charge was neutralized by adding 3 $Na^+$ ions, and the system was supplemented with 0.15 M NaCl. The system was minimized for 1 ns before the simulation. Each system was simulated for 300 ns (in three replicates), generating 1002 frames each, at a temperature of 300 K and a pressure of 1.01325 bar. The system was first equilibrated using the NPT ensemble, with the Nose-Hoover chain thermostat and the Martyna-Tobias-Klein barostat chosen for the simulation. The time step was set to 2.0 fs, and the cutoff method (cutoff radius 9.0 Å) was selected to define the short-range Coulombic interactions. All replicates were randomized, and the trajectories were centred around the substrate. All the molecular visualisations were performed using PyMOL (Schrödinger, LLC. (2020). The PyMOL Molecular Graphics System, Version 2.5).

**Cloning, expression and peptide digestion assay of RufB**

Genomic DNA from *S. atratus* S3_m208_1 was extracted using the Wizard Genomic DNA Purification Kit (Promega) according to the manufacturer's instructions. *E. coli* NEB10ß cells were made chemically competent using the Inoue method (*53*). The *rufB* DNA sequence was codon-optimized for *E. coli* (Thermo Fisher Scientific) and amplified using Q5-Polymerase (NEB). The pET28a-MBP-TEV vector (Addgene) was linearized with PCR. The purified linear vector and insert were mixed at a 1:3 ratio with 15 µL of Gibson assembly master mixture (1X ISO buffer, 10 U/L T5 exonuclease, 2 U/L Phusion polymerase, and 40 U/µL Taq ligase and incubated at 50 °C for 1 h to assemble pET28a-MBP-TEV-RufB. Then, 5 µL of the reaction mix was used to transform chemically competent *E. coli* NEB10b cells, which were plated on LB agar containing 50 µg/mL apramycin. Positive clones containing the plasmid pET28-RufB were screened by colony PCR and confirmed by Sanger sequencing. For expression, a 3 mL inoculum was added to 1 L of LB medium containing 50 µg/mL kanamycin and grown at 180 rpm and 37 °C for ~4 h until the optical density at 600 nm reached 0.6. The culture was then placed on ice for 30 min and induced with 0.05 mM IPTG, followed by incubation at 18 °C for 16 h at 180 rpm. Cells were harvested by centrifugation, and the cell pellet was resuspended in lysis buffer (50 mM Tris, 150 mM NaCl, 10 mM imidazole, 1 mM DTT, pH 7.5), supplemented with 1 mg/mL lysozyme and disrupted by sonication. Cell debris was removed by centrifugation (30 min, 15,000 rcf, 4 °C). The supernatant was incubated with 3 mL of Ni-NTA resin equilibrated with lysis buffer for 1 h at 4 °C, washed with five column volumes of wash buffer I (50 mM Tris, 150 mM NaCl, 20 mM imidazole, pH 7.5) and wash buffer II (wash buffer I with 35 mM imidazole). The protein was eluted with 4 mL of elution buffer (50 mM Tris, 150 mM NaCl, 300 mM imidazole, pH 7.5), concentrated and buffer-exchanged to remove imidazole using a 30 kDa MWCO centrifuge filter unit (Amicon) into storage buffer (50 mM Tris, 150 mM NaCl, 5% glycerol, pH 7.5), and treated with 1.5 U thrombine to 200 µg protein and incubated for 16 h at 4 °C. 0.5 mL of Ni-NTA resin was added to remove impurities MBP-RufB was collected in the flow through fraction and flash-frozen in liquid nitrogen until further use. For the peptide digestion assay, reaction components were combined 100 µL total volume and incubated at 30 °C shaking at 300 rpm with the following final concentrations: RufB (5 µM), **1**-$NO_2$ (100 µM), Tris (50 mM), and NaCl (50 mM). After 16 h, 40 µL assay solution was mixed with 80 µL water +0.1 % FA and 40 µL acetonitrile +0.2 % FA and the protein was removed by centrifugation. For analysis of peptide fragments, 10 µL samples were added to 90 µL $H_2O$ + 0.1% FA and extracted with Pierce C18 (Thermo Fisher Scientific) according to the manufacturer's instruction. 50 µL were injected and analysed on the HR-LCMS. For detection of single amino acids, the 10 µL sample was added to 20 µL water + 0.1% FA and 10 µL acetonitrile and centrifuged for 10 minutes at 15,000 rcf. The supernatant was filtered and analysed by LC-MS using an Agilent 6550 Accurate Mass Q-TOF with ESI-source coupled with an Agilent 1100 HPLC using a Waters Atlantis T3, 5 µm, 4.6 x 50 mm column. The column temperature was 25 °C. MS data were acquired over a range from 50-2500 m/z in positive mode. Auto MS/MS fragmentation was achieved with rising collision energy (35-50 keV over a gradient from 500-2000 m/z) with a frequency of 3 Hz for all ions over a threshold of 500. Elution started with a flow of 0.5 ml/min at 90% $H_2O$ containing 0.1% acetic acid (A) and 10% acetonitrile containing 0.1% acetic acid (B). After 2 min, a linear gradient to 100% B was run for 10 min, followed by an isocratic step with 100% B for 6 min. 20-50 µL were injected per run depending on the concentration of the sample.

**Cloning, Expression and γ$^{18}O_4$-ATP exchange assay of $RufTA_3PCP_3$ /RufH**

DNA sequences of $RufTA_3PCP_3$ and RufH were amplified from genomic DNA with Q5-Polymerase and cloned into pET28 and pRSFDuet-1 (Novagen) for co-expression in a single *E. coli* host using Gibson assembly as described above, to generate pET28- $RufTA_3PCP_3$ and pRSFDuet-1-RufH, which were confirmed for correct assembly by Sanger sequencing. Protein expression and purification was performed analogously to RufB described above. For the γ$^{18}O_4$-ATP exchange assay (*54*) allows for simple and rapid mass spectrometry-based measurement of the activity and substrate specificity of A domains. Three solutions were prepared: Solution 1 with 3 mM γ$^{18}O_4$-ATP and 15 mM $MgCl_2$ in 20 mM Tris (pH 7.5); Solution 2 with 3 mM amino acid and 15 mM Na-pyrophosphate in 20 mM Tris (pH 7.5); and Solution 3 with 5 µM of the enzyme in 5% (v/v) glycerol and 20 mM Tris (pH 7.5). Each solution (2 µL) was mixed and incubated at 22 °C for 90 min. The negative control contained Na-pyrophosphate but no amino acid in Solution 2. The reaction was stopped by adding 6 µL of 9-aminoacridine in acetone (10 mg/mL) and centrifuged at 15,000 x g for 5 min to remove precipitated enzymes. The assay was analyzed by MALDI-TOF mass spectrometry using an AutoFlex III MALDI-Tof/ToF (Bruker) in negative mode, with measurements taken from *m/z* 400-1200 and data analyzed with DataAnalysis 4.2 (Bruker).

Absolute substrate conversion was calculated from the integrals of peaks at m/z 506 for $\gamma^{16}O_4$-ATP and m/z 508, 510, 512, 514 for $\gamma^{18}O_4$-ATP, using the formula absolute substrate conversion = $^{16}O/(^{16}O^{16}+^{18}O)$. Given the equilibrium molar ratio of unlabelled pyrophosphate to $\gamma^{18}O_4$-ATP in the assay is 5:1, an 83.33% detected substrate conversion corresponds to 100% substrate exchange, calculated as % exchange = (absolute substrate conversion) / 0.8333.

#### Deletion mutants of *rufO* and *bytA*

The generation of knock out mutant strains of *S. atratus* S3_m208_1 was performed using homologous recombination and CRISPR-cBEST base editing (*55*). Initially, Cas9 single guide RNA protospacer sequences were selected with CRISPy-web (*56*). The top three sequences were ordered as oligonucleotides with cloning restriction sites and primer sequences for sgRNA amplification. These sgRNAs were PCR-amplified, digested with *Nco*I and *Sna*BI, ligated into similarly digested pCRISPR-cBEST-v2-PermE*, and transformed into *E. coli* NEB10ß. Correct assembly was confirmed by Sanger sequencing, followed by transformation into *E. coli* ET12567 pUZ8002 for conjugation *into S. atratus* S3_m208_1. Apramycin-resistant exconjugants were screened for the stop codon introduction by PCR and confirmed by Sanger sequencing. Due to the small size of *bytA*, the use of CRISPR based systems was not practical, therefore we decided to use the *Streptomyces-E-coli* shuttle vector pYH7 (*57*). After amplification of homologous regions excluding a 12 bp sequence of *bytA*, the fragments were cloned into pYH7 with Gibson assembly (*58*) to yield pYH7_ΔbytA. This plasmid was transformed into *E. coli* NEB10ß and transformed into *Streptomyces atratus S3_m208_1* via conjugation.

#### Feeding experiments

For experiments with deletion mutants *Streptomyces atratus S3_m208_1 rufO*$^{stop}$ and *Streptomyces atratus S3_m208_1 ΔbytA*, a preculture was inoculated from mycelium in TSB and cultivated for 3 d at 30 °C and 180 rpm. Cultivation of main cultures were performed in 5 ml GSS medium (soluble starch 30g/L, glucose 20 g/L, soybean flour 10 g/L, peptone 5g/L, NaCl 3 g/L, $CaCO_3$ 5g/L, corn steep liquor 10 mL) and addition of substrate of interest (0.5 mM final concentration) in 25 mL Erlenmeyer flasks for 7 d at 30 °C and 200 rpm. Cultures were extracted three times with ethyl acetate. Extracts were dissolved in methanol for a final concentration of 2 mg/mL and analyzed with HPLC-MS using a Waters Alliance e2695 Separation Module, equipped with a RP-18 column (Waters XBridge BEH Shield RP18 Column; 3.5 µm, 2.1 x 100mm). A binary solvent mixture of A: 90% acetonitrile + 10% ammonium acetate (5 mM) pH 7.4 and B: 10% acetonitrile + 90% ammonium acetate (5 mM) pH 7.4 was used, with a flow rate of 0.3 mL/min, starting at minute 0-5 with isocratic elution of 10% B, a gradient 10% B → 100% at minute 5-25, and again isocratic conditions of 100% B at minute 25-30.

### Supplementary Tables

**Supplementary Table S1.** Predicted functions of ORFs in the *ruf* BGC in *S. atratus* S3_m208_1 (naming based on the nomenclature by Tomita et al. 2017 (*46*)).

| ORF | Amino acids | Proposed function |
|---|---|---|
| *rufA* | 353 | Streptomycin biosynthesis operon regulator |
| *rufB* | 267 | Serine aminopeptidase |
| *rufC* | 367 | Cytochrome P450 |
| *rufD* | absent | |
| *rufE* | 4819 | Type I polyketide synthase |
| *rufF* | 252 | Thioesterase |
| *rufG* | absent | |
| *rufH* | 80 | MbtH-like protein |
| *rufI* | 380 | Aminotransferase |
| *rufJ* | 161 | Hypothetical protein |
| *rufK* | 359 | ABC transporter |
| *rufL* | 277 | ABC transporter |
| *rufM* | 402 | Cytochrome P450 |
| *bytA* | 5 | RiPP precursor peptide |
| *rufN* | 396 | Nitric oxide synthase |
| *rufO* | 394 | Cytochrome P450 |
| *rufP* | 373 | Tryptophan N-dimethylallyltranferase |
| *rufQ* | 167 | Hypothetical protein |
| *rufR* | 296 | Luciferase-like monooxygenase |
| *rufS* | 399 | Cytochrome P450 |
| *rufT* | 8012 | Nonribosomal peptide synthetase |

**Supplementary Table S2. Comparative analysis of RufNO/IlaMN homologues encoded in various *Streptomyces* and *Nonomuraea* strains.** Identity percentages of homologous RufNO/IlaMN sequences in comparison to the reference sequence of *Streptomyces atratus* ATCC 14046 (*46*). Accession numbers for BGC and P450 enzymes (RufO) are provided alongside the identified precursor peptide sequence for each organism, which is exclusively MRYLH. The asterisk (*) indicates the strain that was used for *in vivo* experiments in this study.

| Strain | Identity [%] | | BGC accession no. | Cyp accession no. | Precursor peptide |
|---|---|---|---|---|---|
| | RufN/IlaM | RufO/IlaN | | | |
| *Streptomyces atratus* S3_m208_1 * | 100 | 100 | - | - | MRYLH |
| *Streptomyces atratus* ATCC 14046 | 96.83 | 97.97 | LC257593.1 | BBA20962.1 | MRYLH |
| *Streptomyces atratus* SCSIO_ZH16 | 97.73 | 97.97 | KY173348.1 | ASX95236.1 | MRYLH |
| *Streptomyces atratus* MJM3502 | 98.23 | 97.72 | CP121214.1 | WPW27215.1 | MRYLH |
| *Streptomyces atratus* 3502J | 97.98 | 97.72 | MN088369.1 | QLF98926.1 | MRYLH |
| *Streptomyces* sp. A108 | 86.62 | 92.93 | JACBYP010000131.1 | MBU6535743.1 | MRYLH |
| *Streptomyces* sp. B4I13 | 87.12 | 92.13 | JAUSZG010000003.1 | MDQ0957474.1 | MRYLH |
| *Streptomyces achromogenes* W4I19-2 | 87.63 | 92.13 | JAUSYA010000001.1 | MDQ0687727.1 | MRYLH |
| *Streptomyces achromogenes* B2I10 | 87.63 | 92.13 | JAUSYH010000001.1 | MDQ0834929.1 | MRYLH |
| *Streptomyces luteosporeus* JCM 4542 | 68.33 | 74.30 | BAAASL010000005.1 | GAA2712158.1 | MRYLH |
| *Nonomuraea solani* CGMCC | 65.90 | 67.18 | FNVT01000022.1 | SEH01838.1 | MRYLH |
| *Nonomuraea basaltis* 160415 | 65.42 | 65.90 | VCJS01000381.1 | TMR88472.1 | MRYLH |
| *Nonomuraea cypriaca* K274 | 64.55 | 65.99 | JADOGI010000057.1 | MBF8188026.1 | MRYLH |

**Supplementary Table S3.** NADH consumption assay data for cytochrome P450 enzymes.

| **Enzyme** | **Substrate** | **Average $K_{cat}$ ($s^{-1}$)** |
|---|---|---|
| $RufO_{SV}$ | Nle-**1** | $0.76 \pm 0.18$ |
| $RufO_{SV}$ + NO | Nle-**1** | $1.47 \pm 0.06$ |
| $RufO_{WT}$ | Nle-**1** | $0.86 \pm 0.06$ |
| $RufO_{WT}$ + NO | Nle-**1** | $1.64 \pm 0.25$ |
| $P450_{Blt}$ | **1** | $3.49 \pm 0.17$ |

**Supplementary Table S4.** Product conversion for cytochrome P450 enzymes with Nle-**1**.

| **Enzyme** | **Cyclised (%)** | **Nitrated Tyrosine (%)** |
|---|---|---|
| $RufO_{WT}$ | 0 | - |
| $RufO_{WT}$ + NO | 0 | $78.1 \pm 8.5$ |
| $RufO_{SV}$ | $19.2 \pm 4.1$ | - |
| $RufO_{SV}$ + NO | $14.9 \pm 3.7$ | $3.3 \pm 1.3$ |
| $P450_{Blt\ WT}$ | 85 | - |
| $P450_{Blt\ WT}$ + NO | 85 | 0 |
| $P450_{Blt\ VP}$ | 0 | - |
| $P450_{Blt\ VP}$ + NO | 0 | 0 |

**Supplementary Table S5.** Selected examples of Cytochrome P450s displaying alternate acid/ alcohol I-helix residues together with their functions. Unusual residues are emboldened in red.

| P450 | Function | Acid | Alcohol | Notes | Ref. |
|---|---|---|---|---|---|
| $P450_{Blt}$ (CYP1251C3) | MRYLH crosslinkiing | Glu | Ser | Tyr-His C-N bond | 59 |
| RufO (CYP1251) | MRYLH Nitration | Glu | **Val** | Tyr nitration | 60 |
| P450cam (CYP101A1) | C-H 5-exo-hydroxylation of (1s)-camphor | Asp | Thr | Archetypal P450 hydroxylase | 61 |
| P450BM3 (CYP102A1) | C-H hydroxylation of fatty acids (mostly ω-1 to ω-3) | Glu | Thr | Fused to redox partner domain | 62 |
| P450SPα (CYP152B1) | α C-H hydroxylation of fatty acids | **Val** | **Ala** | Peroxide driven reaction | 63 |
| OxyB (CYP165B7) | C-O-D crosslinking in teicoplanin biosynthesis | Asp | Asn | X-domain recruitment, PCP-bound substrate | 64 |
| EryF (CYP107A1) | Macrolide hydroxylation of erythromycin | Glu | **Ala** | Substrate-mediated activation | 65 |
| Cyp17A1 | Hydroxylation and C-C bond cleavage of steroids | Glu | Thr | Peroxyanion intermediate for cleavage; cytochrome B5 interaction | 66 |
| AspB | DKP crosslinking during aspergilazine A biosynthesis; | **Pro** | Thr | Ferric superoxide intermediate | 67 |
| CypX (CYP134A1) | DKP aromatisation and N-oxidation | Glu | **Pro** | Multiple step oxidation | 68 |
| XplA (CYP177A1) | Reductive denitration of RDX | **Met** | **Ala** | Reductive transformation | 69 |
| TxtE | Nitration of Trp in DKP biosynthesis of thaxtomin A | **Pro** | Thr | Uses NO + postulated ferric superoxide intermediate | 70 |
| Sas16 (CYP) | Desaturation of Tyr in acyl-Thr-Tyr dipeptide in WS9326A biosynthesis | Glu | **Phe** | PCP-bound substrate | 71 |
| CYP79A1 | Oxime formation from Tyr | Asp | Asn | Multiple step oxidation | 72 |
| Cyp74A1 | Allene oxide synthase | **Gly** | **Gly** | Rearrangement of fatty acid peroxide | 73 |

**Supplementary Table S6.** Primers used in this study.

| Primer | Sequence |
|---|---|
| rufO_V240S_fwd | CGAGTCCGTGTCCCGATTCCTCGGC |
| rufO_V240S_rev | CTCGAAGCCTGCATGAAAAAACGCTC |
| rufO_V240SP241V_fwd | CGAGTCCGTGTCCCGATTCCTCGGC |
| rufO_V240SP241V_rev | GGACACGGACTCGAAGCCTGCATG |
| P450Blt_S239V_fwd | ACGAGGTGGTGTCCCGTCTGCTGGG |
| P450Blt_S239V_rev | GACACGTGCTCGTAACCAGCGTGCA |
| P450Blt_S239VV240P_fwd | CGAGGTGCCGTCCCGTCTGCTGGGC |
| P450Blt_S239VV240P_rev | CGGGACGGCACCTCGTAACCAGCGTGCAG |
| pYH7_ ΔbytA_up_for | GCTTGCGGCAGCGTGAAGCTGATCTTCCGGAAGTCGGTG |
| pYH7_ ΔbytA_up_rev | TGAAAGTAGAAACGAGACC |
| pYH7_ ΔbytA_down_for | TGGTCTCGTTTCTACTTTCACATATCGATTCACCTCCCCTG |
| pYH7_ ΔbytA_down_rev | GACCTGCAGGCATGCAAGCTCACCGCATCGGAAATCGTC |
| pYH7_ ΔbytA_screen_for | CGTTGTAGGCCAGCCGCAG |
| pYH7_ ΔbytA_screen_rev | TCGAGGAGCTGATGCGCTG |
| rufO_sgRNA1_for | CATGCCATGGGGCGTGCCAGTAGACCGGCT GTTTTAGAGCTAGAAATAGC |
| rufO_sgRNA2_for | CATGCCATGGCGGCGTGCCAGTAGACCGGC GTTTTAGAGCTAGAAATAGC |
| rufO_sgRNA3_for | CATGCCATGGCATCAGCCAGGAGTCGAGGC GTTTTAGAGCTAGAAATAGC |
| sgRNA_rev | ACGCCTACGTAAAAAAAGCACCGACTCGGTGCC |
| RufB_MBP_TEV_for | CTGAGAACCTGTACTTCCAATCCATGGATTTTCCGACACGTC |
| RufB_MBP_TEV_rev | TGCTGTCCACCAGTCATGCTAGCTTACGGAGCATCCAGAAATGC |
| RufB_codonopti _for | TGGTGCCGCGCGGCAGCCATATGGATTTTCCGACACGTCAGG |
| RufB_codonopti _rev | GAGTGCGGCCGCAAGCTTGGTTACGGAGCATCCAGAAATGCAC G |
| pet28a_linearized_for | CCAAGCTTGCGGCCGC |
| pet28a_RufB_linearized_re v | ATGGCTGCCGCGCGG |
| rufB_screen_for | CCATGAACGCTCGGAACG |
| rufB_screen_rev | TCAGGGGGCGTCGAGG |
| rufTA3PCP3_for | TAGAAGCTTACCCCTCCGACAGGTGCCG |
| rufTA3PCP3_for | TATCATATGCTGCTTGCCCCGCAGGAGCG |
| rufN/O_screen_for | GTGAGTTCTTTCCGCAGG |
| ΔbytA_screen_for | TTTCTACTTTCACATATCG |
| rufM_screen_for | CGTACCGCTCAGCCTGC |
| rufM_screen_rev | CCAGCTCGGTGTCGGTG |

### Supplementary Figures

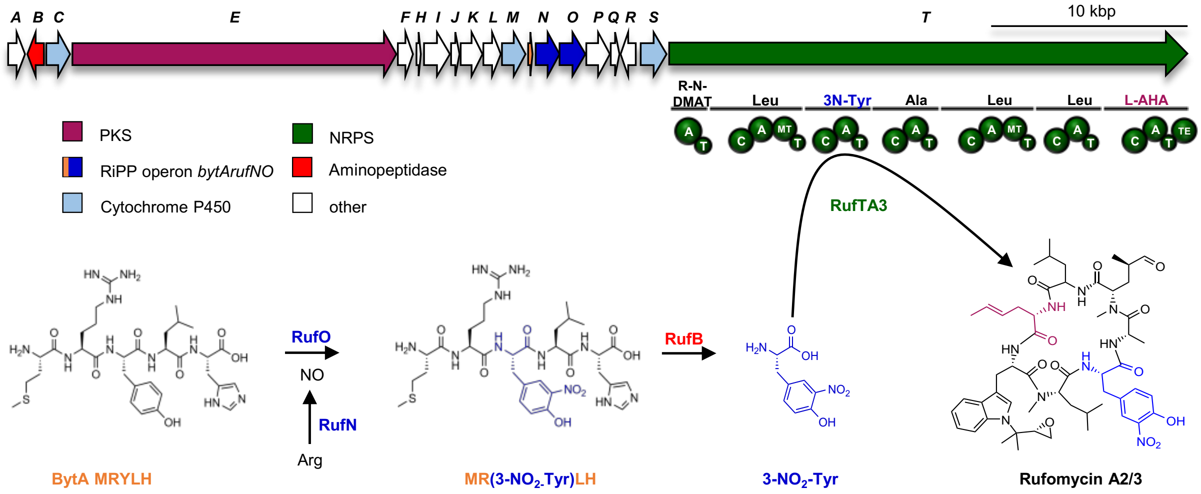

**Supplementary Figure S1. The rufomycin BGC harbors a unique pathway for 3-$NO_2$-Tyr biosynthesis.** Schematic overview on the *ruf* BGC from *S. atratus* S3_m208_1 and 3-$NO_2$-Tyr biosynthesis and incorporation.

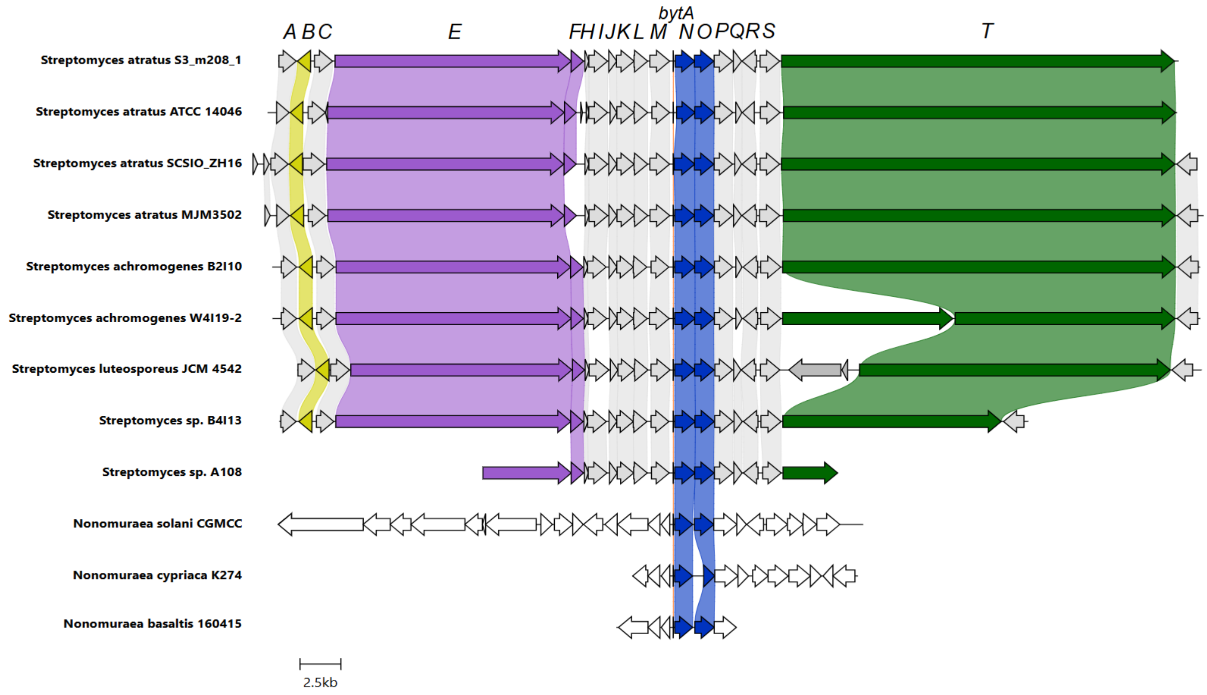

**Supplementary Figure S2. The biarylitide precursor gene *bytA* is present in all detected homologous rufomycin/ilamycin gene clusters.** Comparison of gene clusters harbouring the *rufNO* operon and the biarylitide precursor peptide. Summary phylogeny of rufomycin/ilamycin BGCs. All clusters encode the precursor peptide MRYLH directly upstream of *rufNO*. Homologous genes in each gene cluster are linked. Figure to scale.

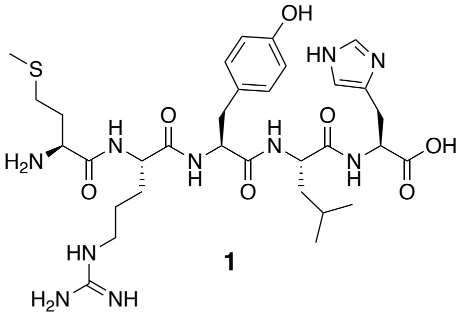

**1**

Chemical Formula: $C_{32}H_{50}N_{10}O_7S$
Exact Mass: 718.35846

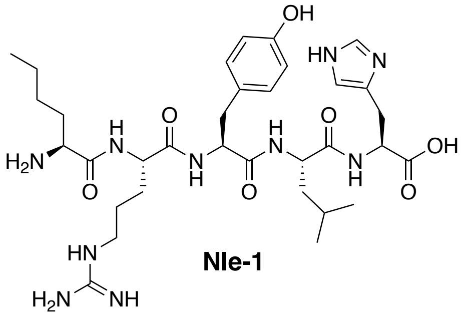

**Nle-1**

Chemical Formula: $C_{33}H_{52}N_{10}O_7$
Exact Mass: 700.40204

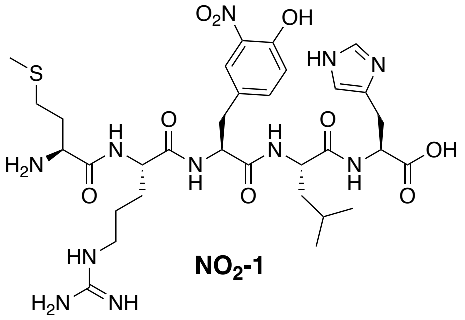

**$NO_2$-1**

Chemical Formula: $C_{32}H_{49}N_{11}O_9S$
Exact Mass: 763.34354

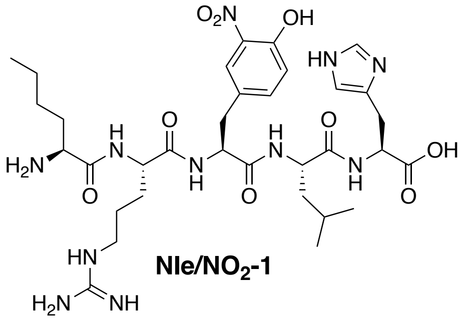

**Nle/$NO_2$-1**

Chemical Formula: $C_{33}H_{51}N_{11}O_9$
Exact Mass: 745.38712

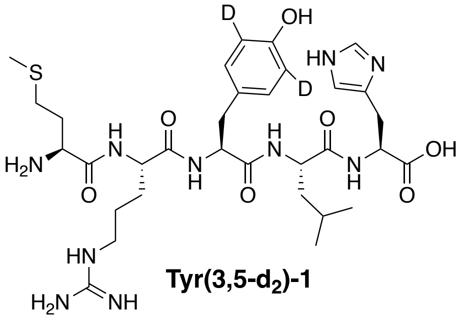

**Tyr(3,5-$d_2$)-1**

Chemical Formula: $C_{32}H_{48}D_2N_{10}O_7S$
Exact Mass: 720.37102

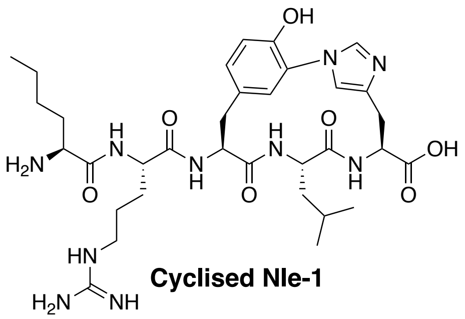

**Cyclised Nle-1**

Chemical Formula: $C_{33}H_{50}N_{10}O_7$
Exact Mass: 698.38639

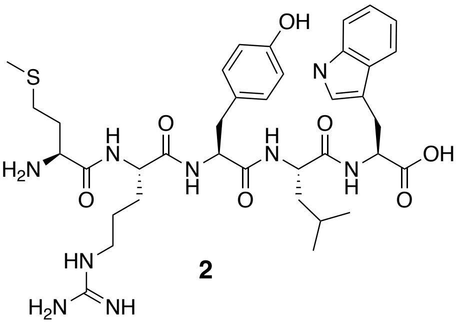

**2**

Chemical Formula: $C_{37}H_{52}N_9O_7S$
Exact Mass: 766.37104

**Supplementary Figure S3.** Structures of the peptides used in this study. Syntheses/characterisation of **1**,(51) Nle-**1**,(51) Tyr-(3,5-$d_2$)-**1**,(*50*) **2**,(*51*) and characterisation of enzyme Cyclised Nle-**1** (*59*) as previously reported.

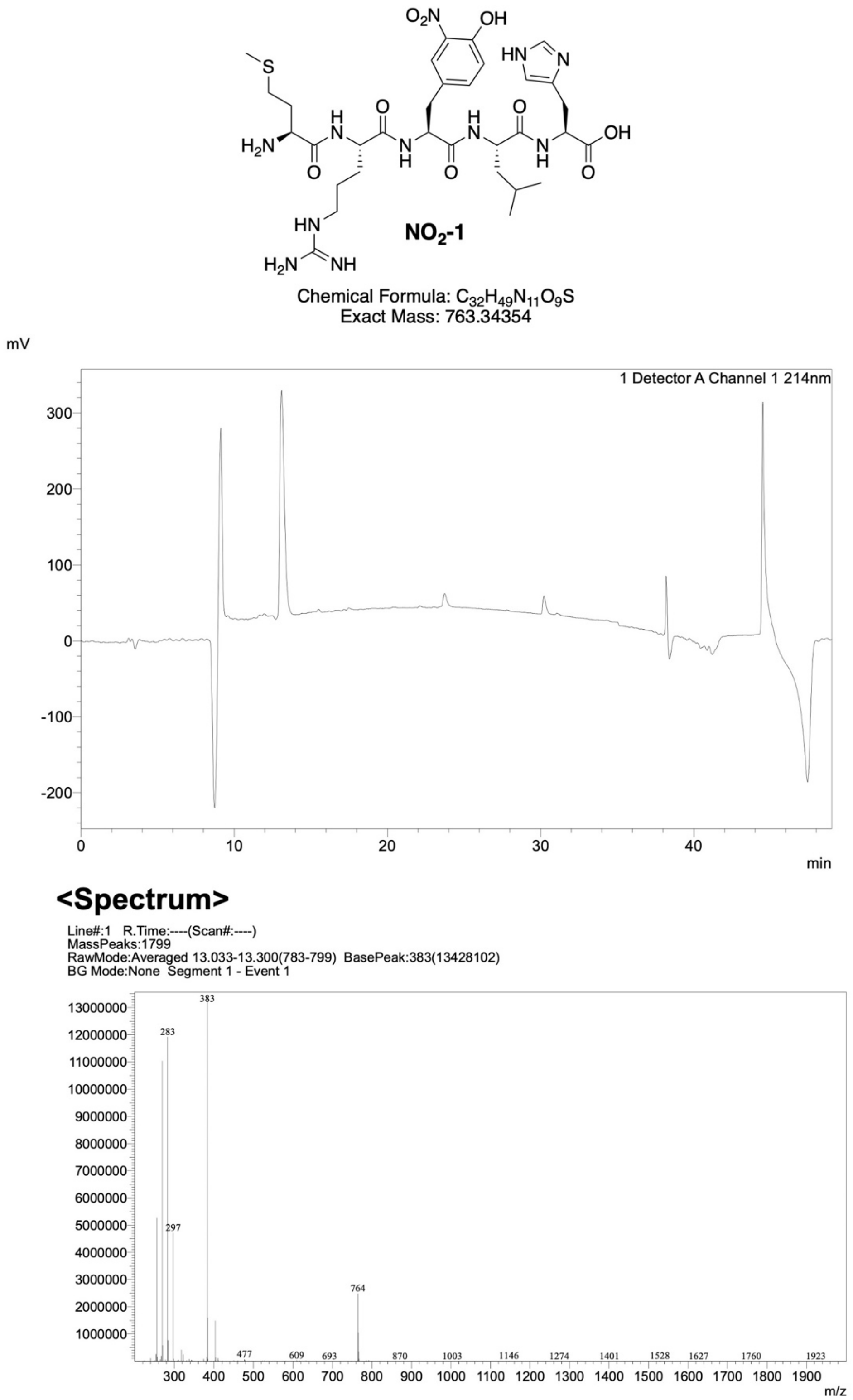

**Supplementary Figure S4.** LC and MS analysis of synthetic peptide $NO_2$-**1**.

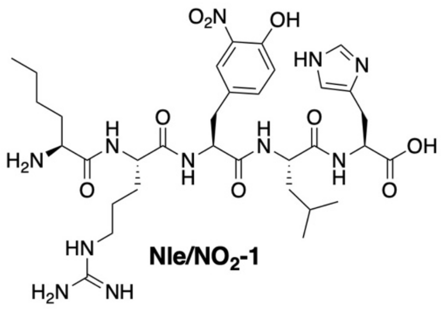

Chemical Formula: $C_{33}H_{51}N_{11}O_9$
Exact Mass: 745.38712

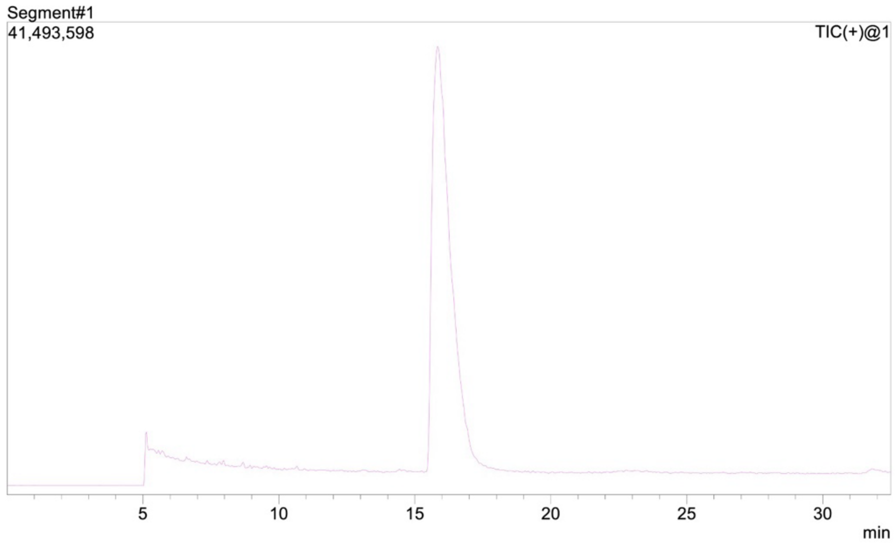

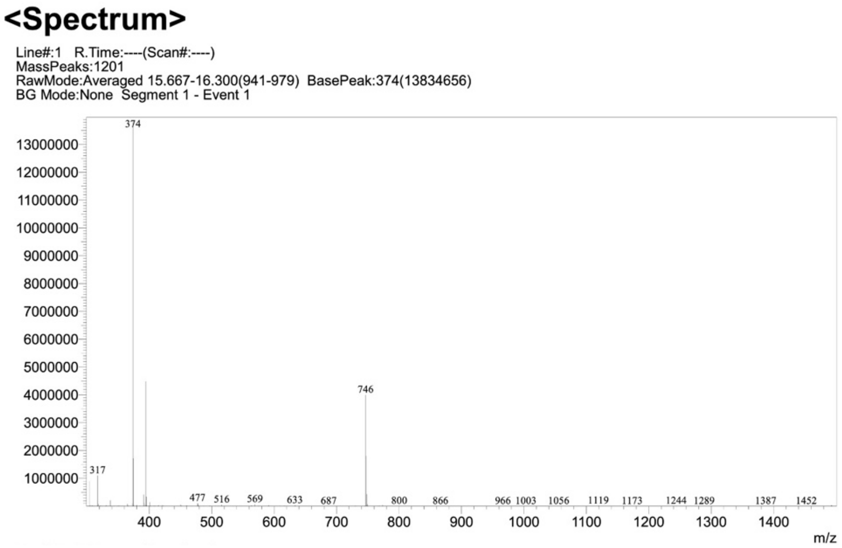

**Supplementary Figure S5.** TIC and MS analysis of synthetic peptide Nle/$NO_2$-**1**.

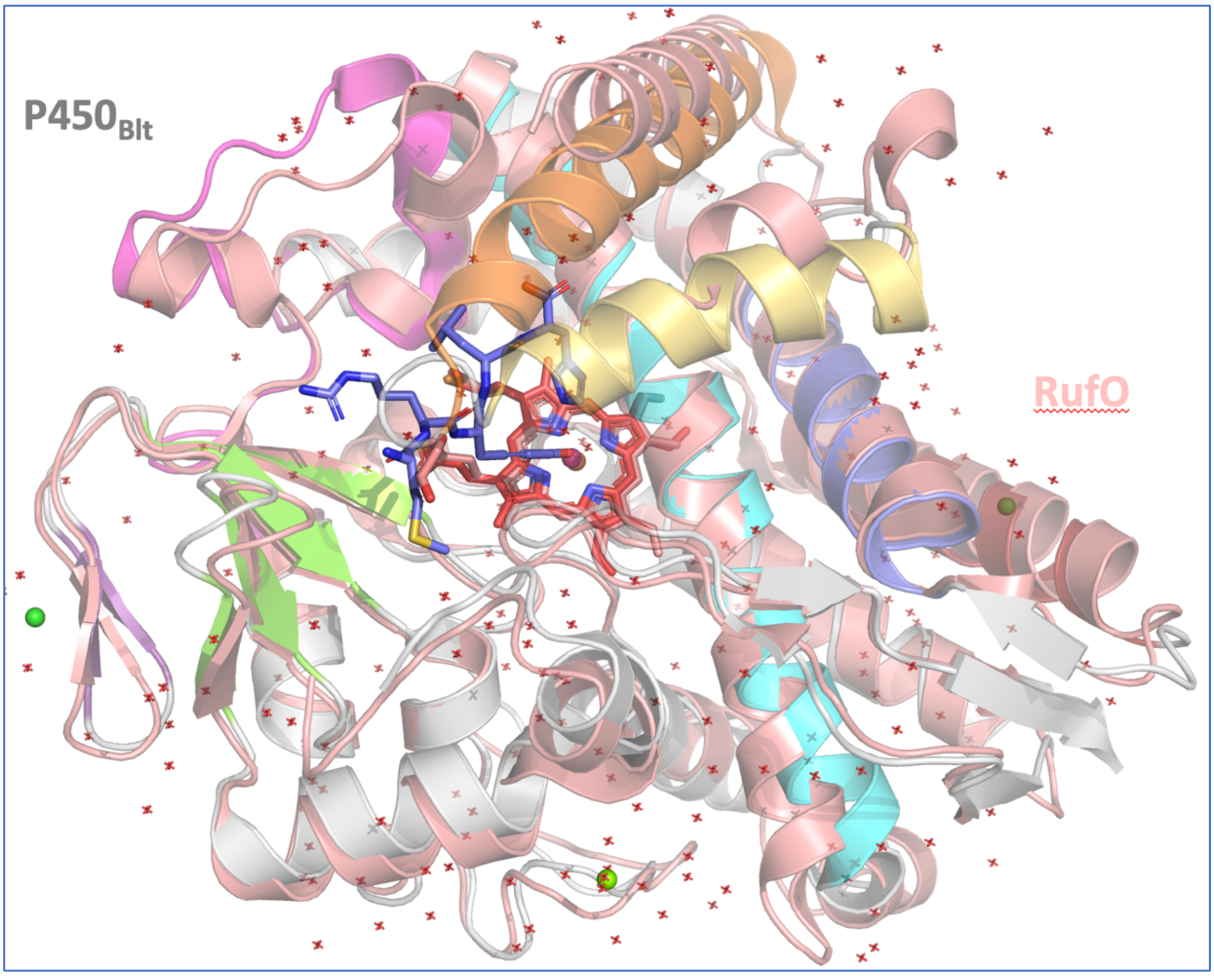

**Supplementary Figure S6.** Structure of RufO (PDB: 8SPC, chain A) (*74*) overlayed on the structure of $P450_{Blt}$ (PDB: 8U2M, chain A) (*59*). RufO – salmon; $P450_{Blt}$ – grey, heme – red sticks, MRYLH peptide substrate – blue sticks. Specific $P450_{Blt}$ secondary structure elements coloured (B/C loop – pink, D-helix – brick red, E-helix – slate blue, F-helix – yellow, G-helix – orange, I-helix – turquoise, $\beta_1$-sheet – green, $\beta_2$-sheet – purple).

```
P450Blt   MADEATAAVRDPIYDPLAPSVIADPYPFYRKLRETNTVHWHEFLDSWVVTGYAECRQVLG 60
RufO      MTVSPAPEHTDPLFSPLDPAVLADPYPVYRRLRETHPVYWHAGLDSWLMTRHADCTAILR 60
          *: . :    **::.** *:*:*****.**:****: *:**  ****::* :*:*  :*

P450Blt   DTTNFGSDFRRIDVEIPDTQLSVQSLDPPEHGAIRHLLVSALHEQPLSTVRQQFAAIAAQ 120
RufO      DPGRFSTDFRKIDIPTPPTLLSLQTLDPPDQTPLRHLALDAVRAQDLDALRKELTLFADQ 120
          *  .*.:***:**:  * * **:*:****::  :*** :.*:: * *.::*:::: :* *

P450Blt   HLAELSGQPGTVDLVSRFARPVALRTITAFLGVPPPD-GAGFEQWSNAIVRSMDAGIEPA 179
RufO      LLDELAD-RESFDFIHDYADVFTLRAITRFIGVEPPETDEAFARFNDDLDHSMDAQLDPD 179
           * **:.   :.*::  :*  .:**:** *:** **: . .* ::.: : :**** ::*

P450Blt   RAEPGNQARAELSRLVTHWLAEADERGFVGAARRAARAQD-VPAAVLANSLRAVLHAGYE 238
RufO      AEEPGLRARAHFNDLVRSWLGDPGPHGVLPDVVRLLPGSGVEADDVLVNSVRAFFHAGFE 239
            *** :***.:. **  **.: . :*.:  . *   ...     **.**:**.:***:*

P450Blt   SVSRLLGGVLARLVRHPELLAGPATRD-ADEALVDELIRLDGPVQADARVCVRDQPVGAQ 297
RufO      VPSRFLGNALAALLATPGAWEQLVRGDVGLDTAVEELIRYVGPVQALARACLQDTELGGM 299
            **:**..** *:  *      .  * . :: *:****  ***** **.*::*  :*.

P450Blt   LVRRGDVLVLFIAAANRDPAVFPDPDAVRLTRRRGLHLAFGRGAHACLGAGLATLQLREV 357
RufO      AVKEGQVVTALIGAANRDPDQFPDPETLRLDRKPNNHLGFGRGAHSCLGLNVARIEAHVT 359
           *:.*:*:. :*.******  ****:::** *: . **.******:*** .:* :: : .

P450Blt   LGALRAGGLRLAPAGPAAYEPTATLRGLAELPVSVRQPHRTD    399
RufO      LGALLRH-PGVRSAGEPVVRPNGTLRGLSRLPLTLG------    394
          ****      :  **  . .*..*****:.**:::
```

**Supplementary Figure S7.** Alignment of RufO (*46*) and P450$_{Blt}$ (*59*) with key mutations in the I-helix coloured red and highlighted in yellow. Additional residue possibly contributing alterations in RufO I-helix rotamers (PDB: 8SPC, chain A) is highlighted in yellow, although incomplete dentistry does not allow this to be unambiguously assigned. Lack of density in the F/G-helices of the RufO structure are highlighted in grey.

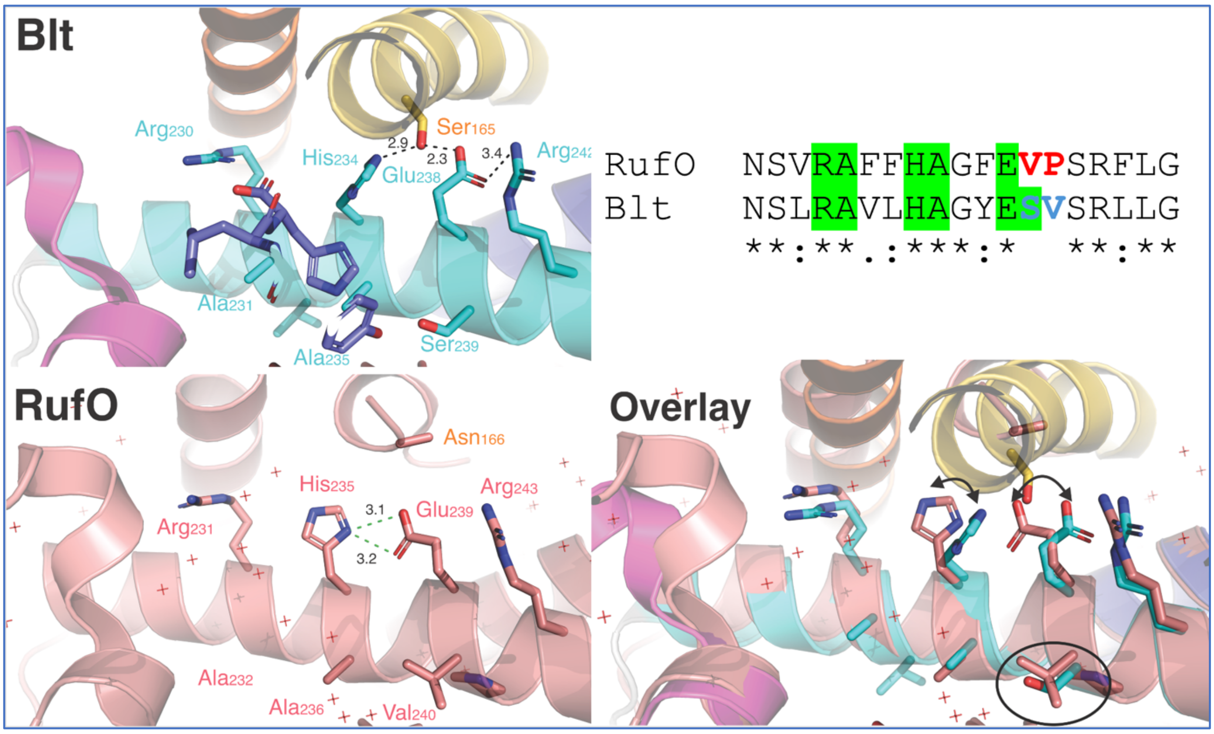

**Supplementary Figure S8.** Structural comparison of the active site I-helix residues of $P450_{Blt}$ (upper left; PDB: 8U2M, chain A) (*59*) and RufO (lower left; PDB: 8SPC, chain A) (*75*); the colour scheme is the same as in the previous figure, residues are labelled and H-bonding distances shown in Angstrom. Comparison of the key I-helix residues via alignment (upper right) and overlay of the $P450_{Blt}$ and RufO structures with differences highlighted (lower right).

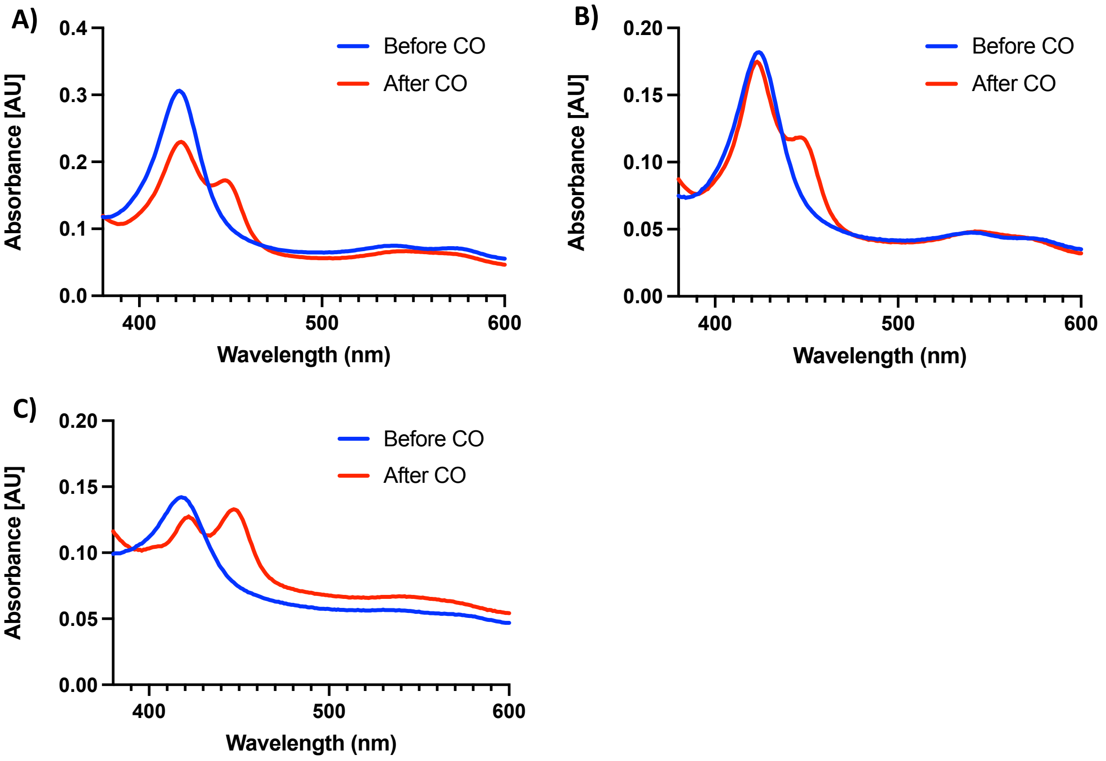

**Supplementary Figure S9.** CO difference spectra of the P450s in this study; **A)** $RufO_{WT}$, **B)** $RufO_{SV}$, **C)** $P450Blt_{VP}$. $P450Blt_{WT}$ as previously published. (*51*)

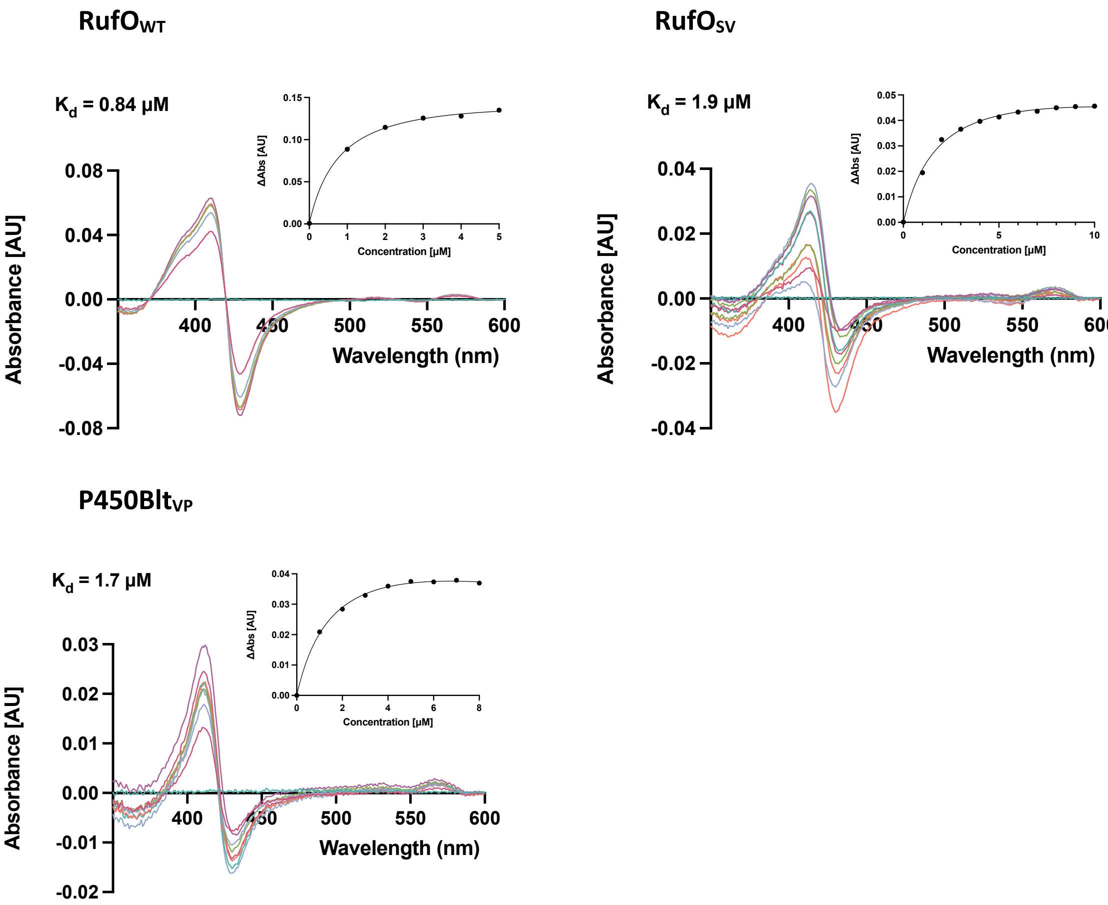

**Supplementary Figure S10.** Titration of the cytochrome P450s RufO, RufO$_{SV}$ and P450Blt$_{VP}$, with Nle-**1** $K_d$ calculated as described in methods. Binding to P450$_{Blt}$ as previously published. (50)

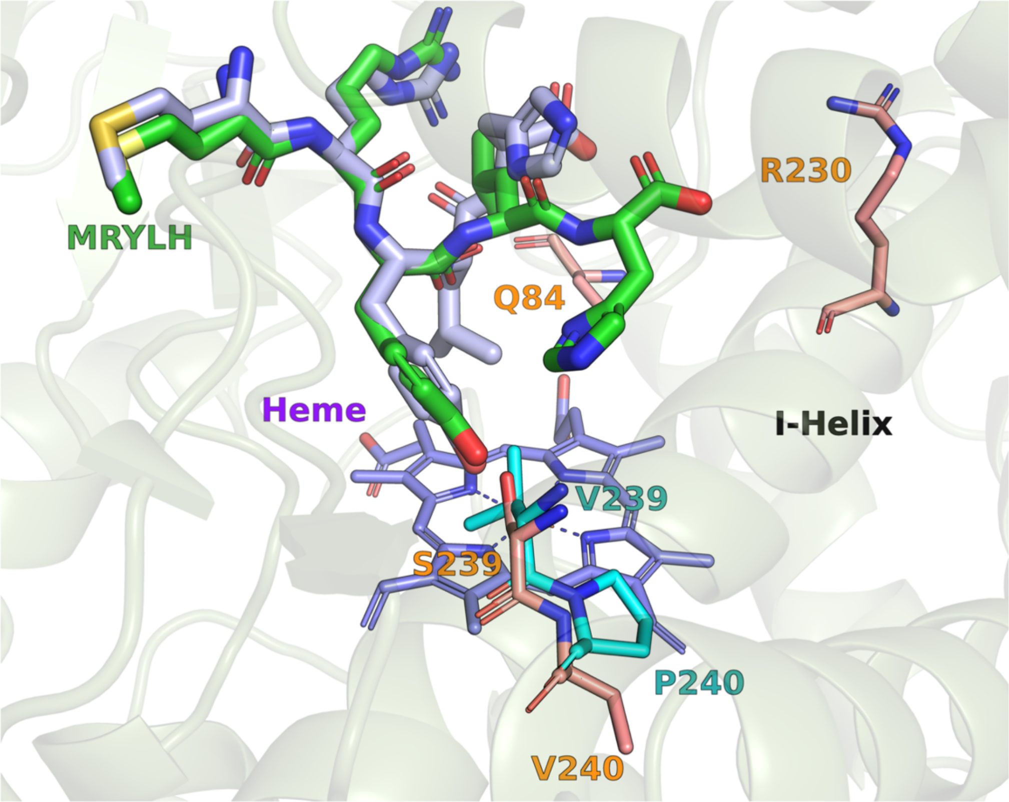

**Supplementary Figure S11.** Overlapping binding orientations of representative poses from molecular dynamics simulations of MRYLH peptide **1** bound in the active site of $P450_{Blt}$ (green) and the Ser239Val/Val240Pro variant (grey) after 300 nanoseconds. Mutated residues Val239 and P240 are shown in cyan, while wild-type residues are shown in orange. Note the major shifts in position of the Leu-4 and His-5 residues of **1**.

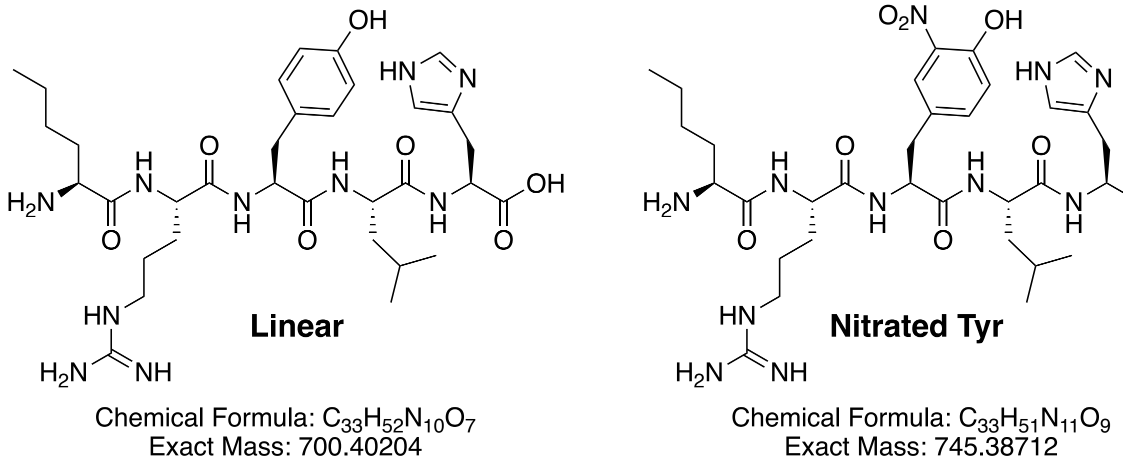

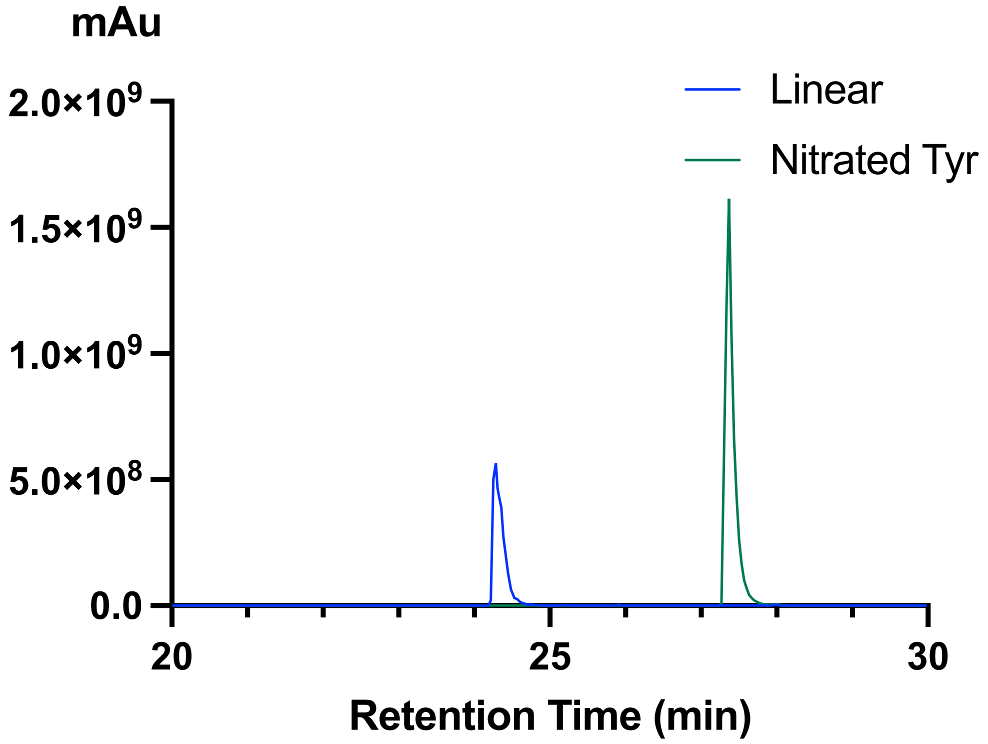

**Supplementary Figure S12.** Nitration data for Nle-**1** using RufO$_{WT}$ in the presence of nitric oxide. The structures of linear and nitrated tyrosine-containing peptide are shown. Overlapping ESI traces for linear and nitrated tyrosine-containing peptides based on HRMS analysis.

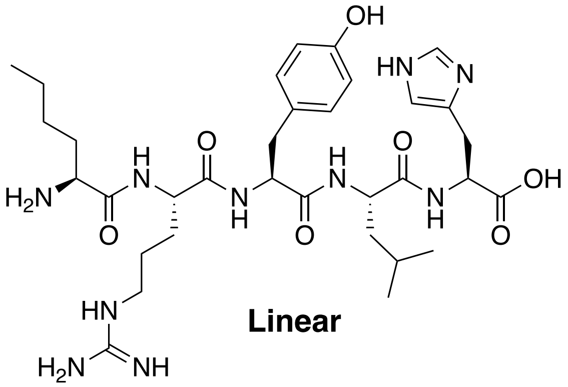

Chemical Formula: $C_{33}H_{52}N_{10}O_7$
Exact Mass: 700.40204

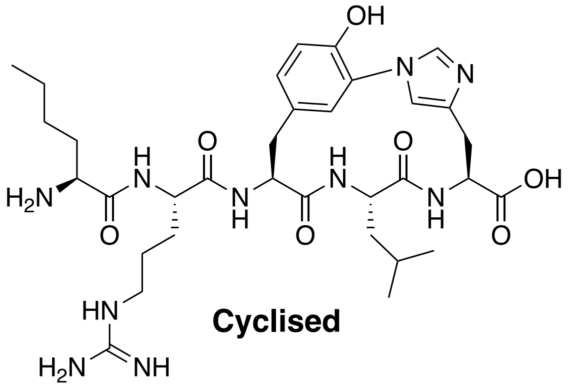

Chemical Formula: $C_{33}H_{50}N_{10}O_7$
Exact Mass: 698.38639

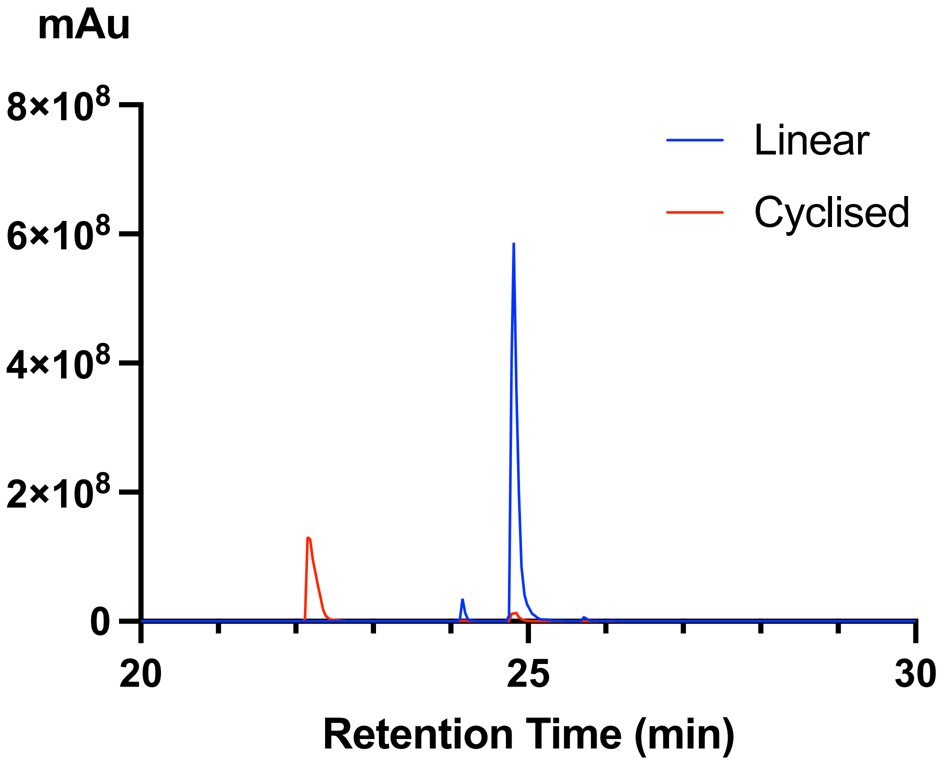

**Supplementary Figure S13.** Cyclisation data for Nle-**1** using $RufO_{SV}$ in the absence of nitric oxide. The structures of linear and cyclised peptide are shown. Overlapping ESI traces for linear and cyclised peptides based on HRMS analysis.

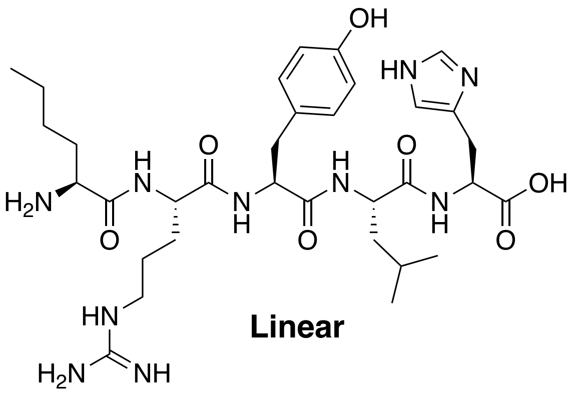

**Linear**

Chemical Formula: $C_{33}H_{52}N_{10}O_7$
Exact Mass: 700.40204

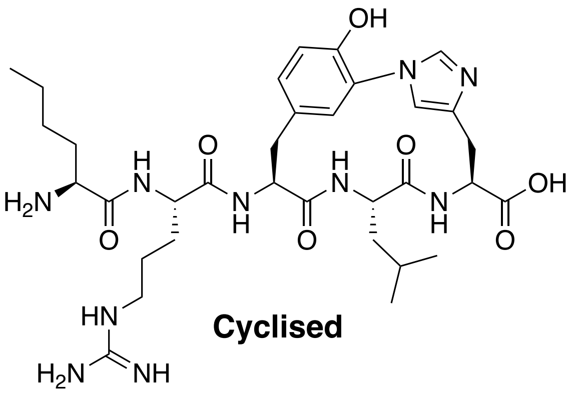

**Cyclised**

Chemical Formula: $C_{33}H_{50}N_{10}O_7$
Exact Mass: 698.38639

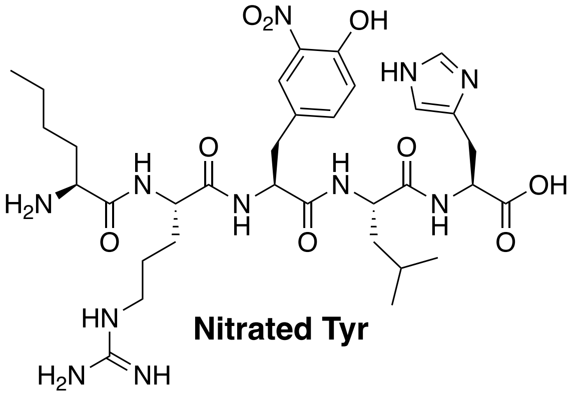

**Nitrated Tyr**

Chemical Formula: $C_{33}H_{51}N_{11}O_9$
Exact Mass: 745.38712

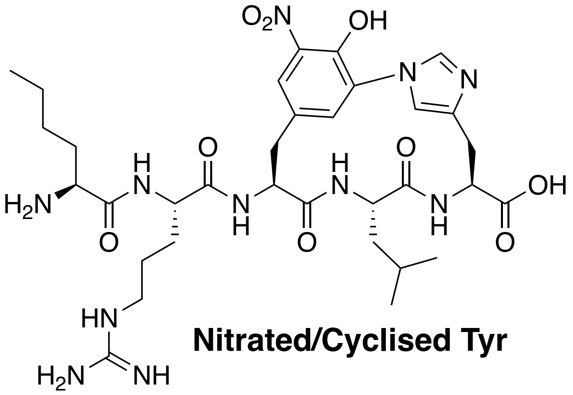

**Nitrated/Cyclised Tyr**

Chemical Formula: $C_{33}H_{49}N_{11}O_9$
Exact Mass: 743.37147

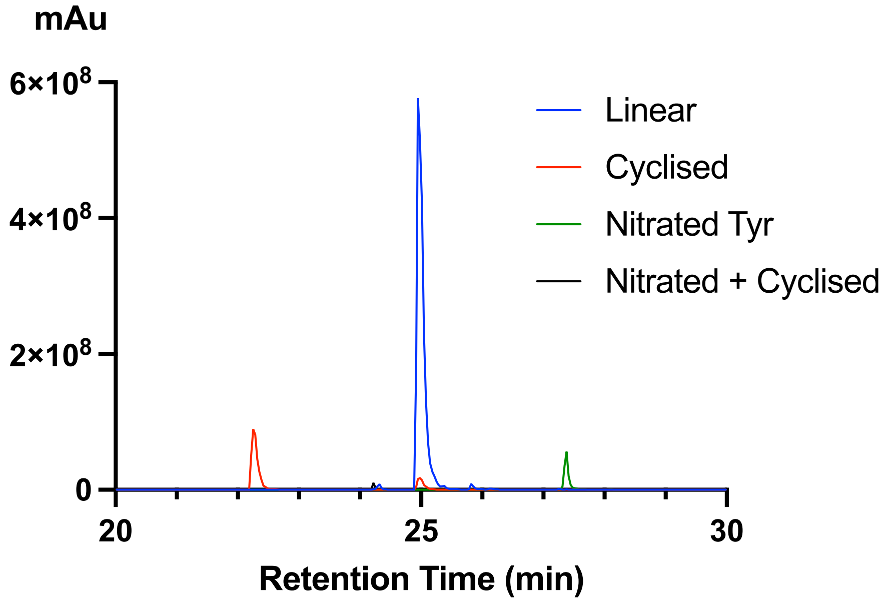

**Supplementary Figure S14.** Cyclisation data for Nle-**1** using $RufO_{SV}$ in the presence of nitric oxide. The structures of linear, cyclic, nitrated tyrosine-containing peptide and nitrated tyrosine and cyclised peptides are shown. Overlapping ESI traces for linear, cyclic, nitrated tyrosine-containing peptide and nitrated tyrosine and cyclised peptides based on HRMS analysis.

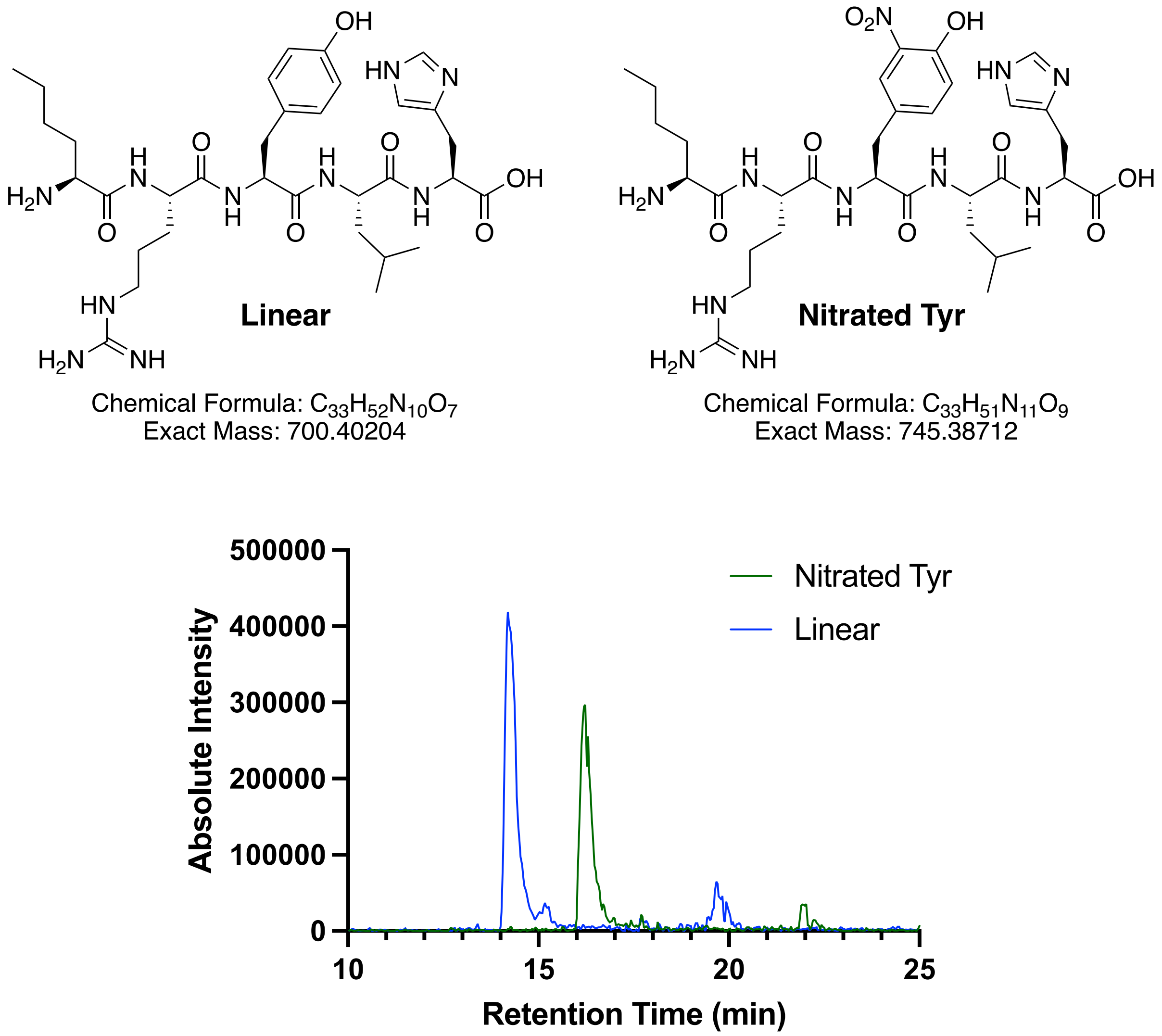

**Supplementary Figure S15.** Nitration data for Nle-**1** using $RufO_{WT}$ in the presence of nitric oxide with sodium dithionite and no redox partners. The structures of linear and nitrated tyrosine-containing peptide are shown. Overlapping ESI traces for linear and nitrated tyrosine-containing peptides based on LCMS analysis.

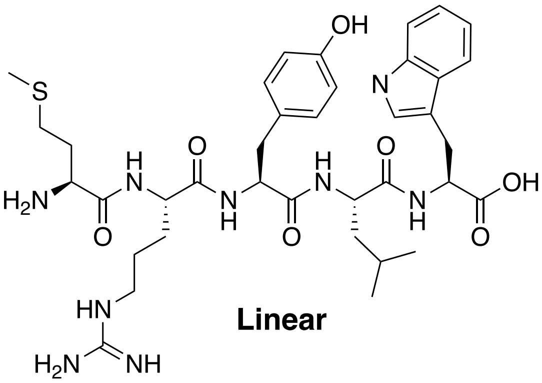

**Linear**

Chemical Formula: $C_{37}H_{52}N_9O_7S$
Exact Mass: 766.37104

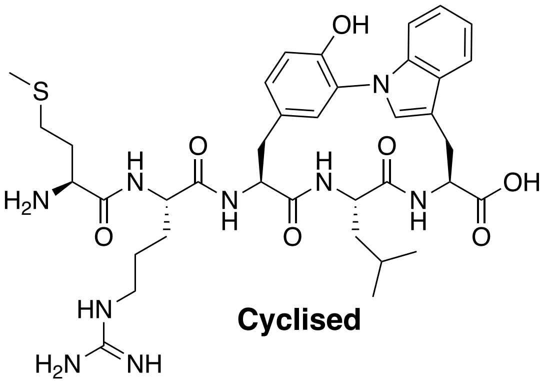

**Cyclised**

Chemical Formula: $C_{37}H_{51}N_9O_7S$
Exact Mass: 765.36322

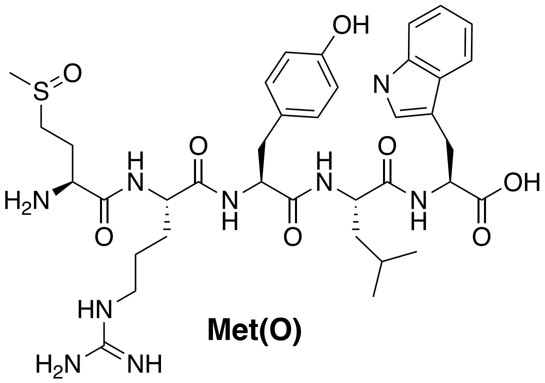

**Met(O)**

Chemical Formula: $C_{37}H_{52}N_9O_8S$
Exact Mass: 782.36596

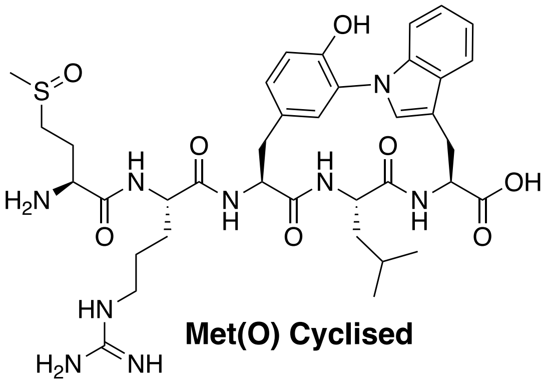

**Met(O) Cyclised**

Chemical Formula: $C_{37}H_{51}N_9O_8S$
Exact Mass: 781.35813

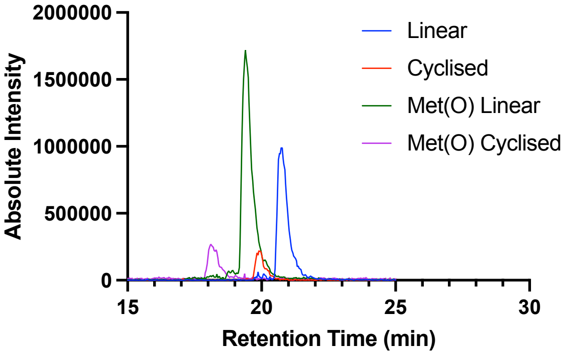

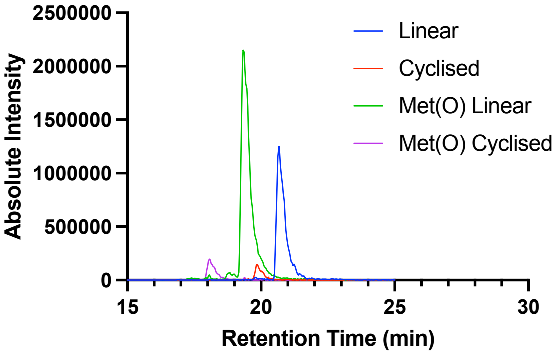

**Supplementary Figure S16.** Cyclisation data for **2** using $RufO_{WT}$ (left) and $RufO_{SV}$ (right). The structures of linear and cyclised peptides along with the oxidised methionine versions of these peptides are shown. Overlapping ESI traces for the peptides based on LCMS analysis.

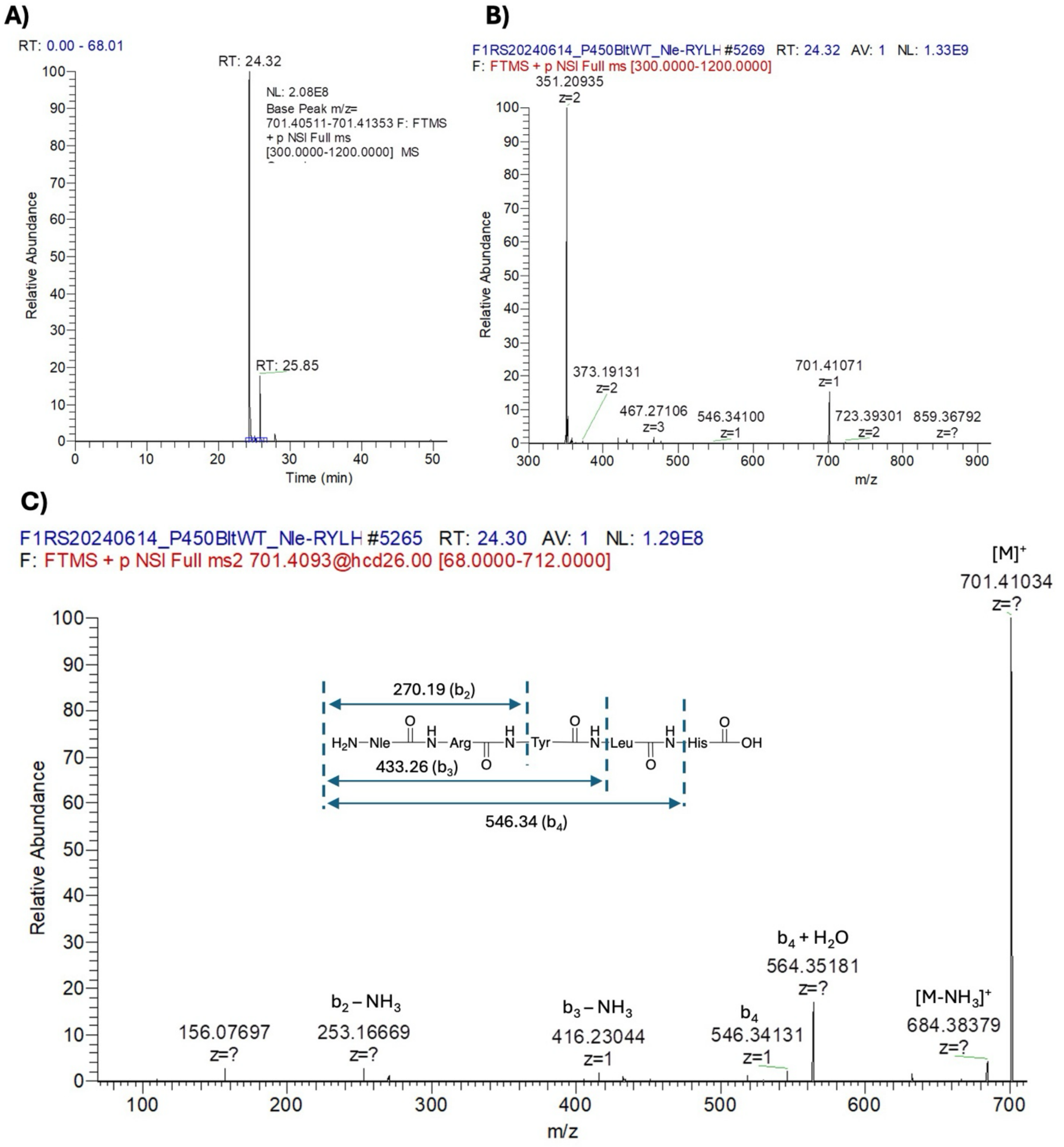

**Supplementary Figure S17.** HRMS and $MS^2$ data for linear peptide Nle-**1**. **A)** EIC trace of Nle-**1**. **B)** HRMS spectrum for Nle-**1**. **C)** Annotated $MS^2$ for Nle-**1**.

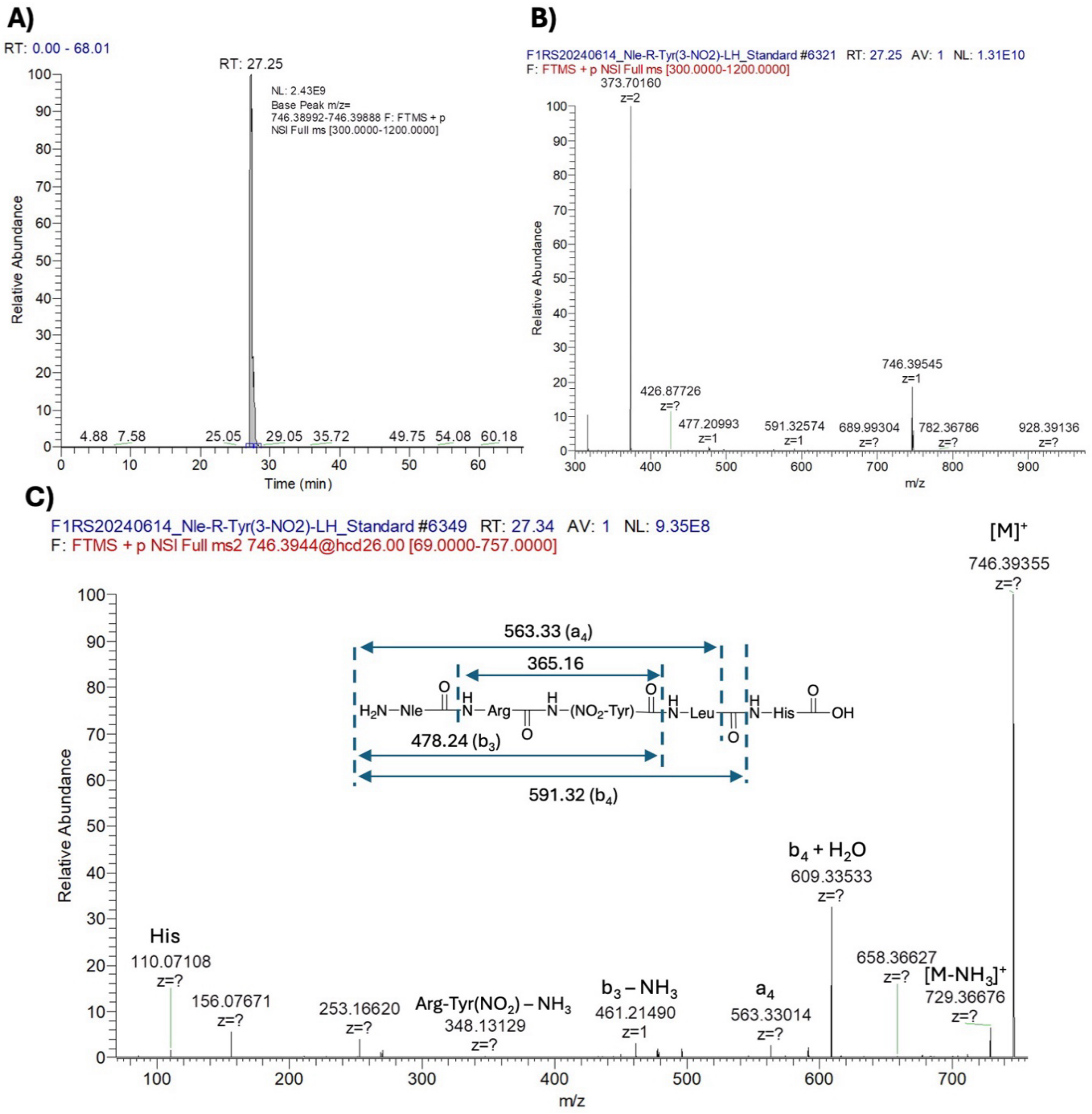

**Supplementary Figure S18.** HRMS and $MS^2$ data for nitrated tyrosine-containing peptide Nle/$NO_2$-**1**. **A)** EIC trace of Nle/$NO_2$-**1**. **B)** HRMS spectrum for Nle/$NO_2$-**1**. **C)** Annotated $MS^2$ for Nle/$NO_2$-**1**.

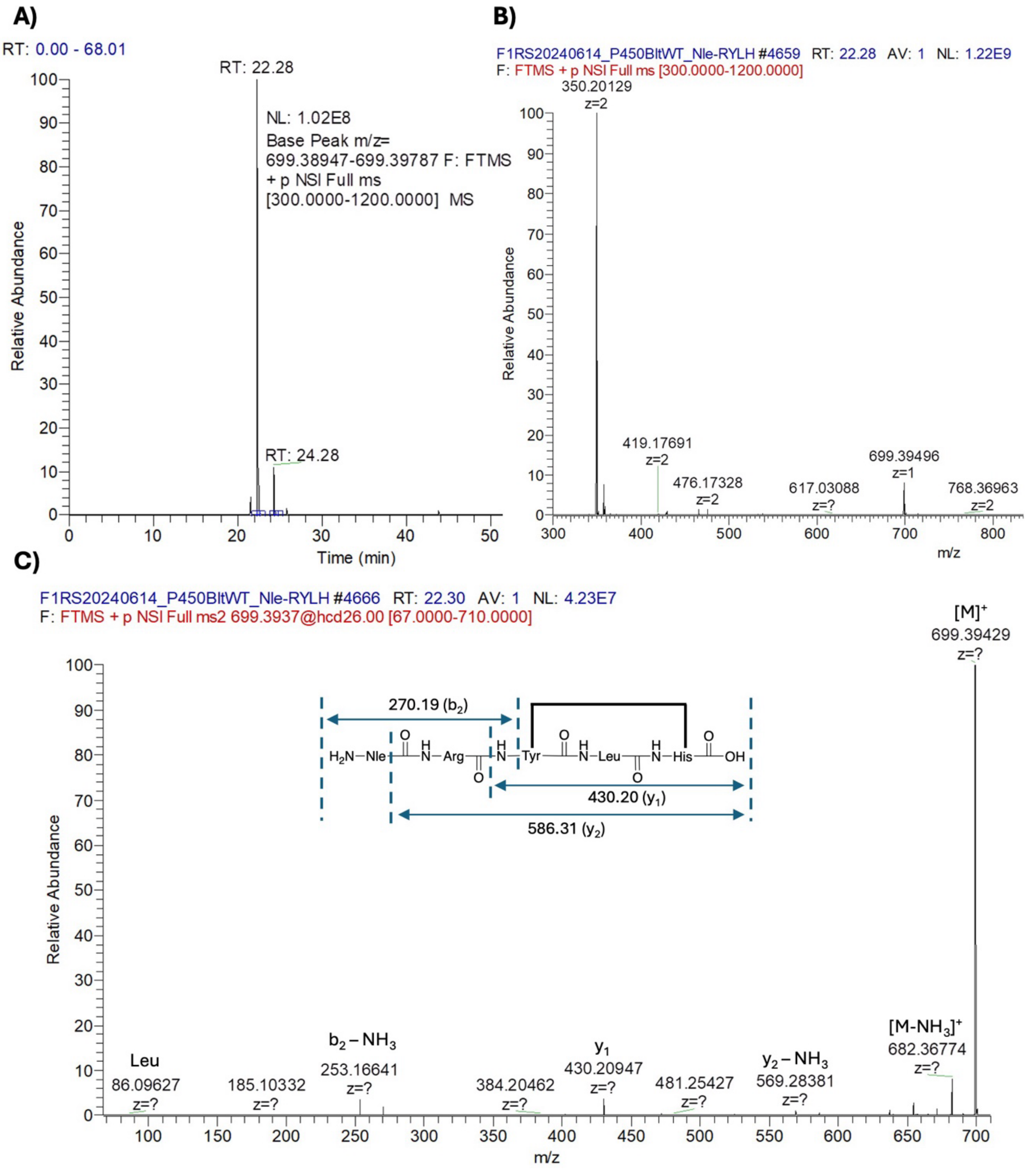

**Supplementary Figure S19.** HRMS and MS$^2$ data for cyclic peptide Cyclised Nle-**1**. **A)** EIC trace of Cyclised Nle-**1**. **B)** HRMS spectrum for Cyclised Nle-**1**. **C)** Annotated MS$^2$ for Cyclised Nle-**1**.

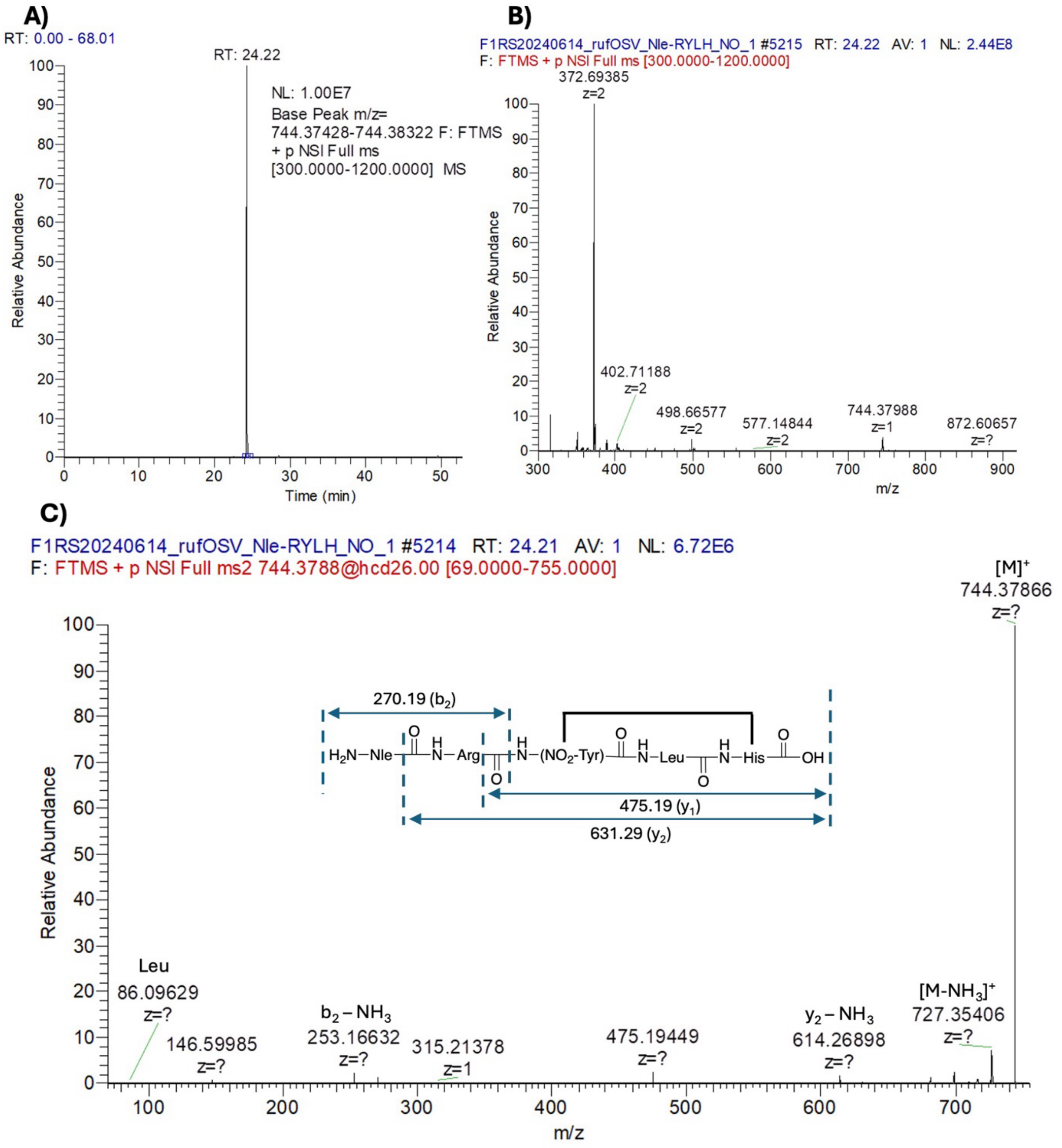

**Supplementary Figure S20.** HRMS and $MS^2$ data for cyclised and nitrated tyrosine-containing peptide Nle/$NO_2$-**1**. **A)** EIC trace of Cyclised Nle/$NO_2$-**1**. **B)** HRMS spectrum for Cyclised Nle/$NO_2$-**1**. **C)** Annotated $MS^2$ for Cyclised Nle/$NO_2$-**1**.

| **Protein (*75*)** | **$A_6$ selectivity code (*76*)** | | | | | | | | | | **Description** |
|---|---|---|---|---|---|---|---|---|---|---|---|
| $A_6$ Teicoplanin | D | A | S | T | I | A | G | V | C | K | Tyr permissive pocket (teicoplanin-type) |
| $A_6$ UK-68597 | D | A | S | T | I | A | G | V | C | K | |
| $A_6$ Balhimycin | D | A | S | T | L | G | A | I | C | K | Tyr permissive pocket (balhimycin-type) |
| $A_6$ Vancomycin | D | A | S | T | L | G | A | I | C | K | |
| $A_6$ Chloroerymomycin | D | A | S | T | L | G | A | I | C | K | |
| $A_6$ Arylomycin | D | A | S | T | V | A | A | V | C | K | Tyr specific pocket |
| $A_6$ A40926 | D | A | S | T | V | A | A | V | C | K | |
| $A_6$ Complestatin | D | A | S | T | V | A | A | V | C | K | |
| $A_6$ Kistamicin | D | A | S | T | V | A | A | V | C | K | |
| $A_6$ Kistamicin-mut | D | A | S | T | I | A | G | V | C | K | Tyr permissive pocket |
| $A_3$ Rufomycin (*46*) | D | A | S | T | A | A | A | V | G | K | $NO_2$-Tyr pocket |

**Supplementary Figure S21.** Comparative analysis of the RufT $A_3$ domain specificity-conferring code with other notable tyrosine activating pockets. These include those with tolerance for β-hydroxylation and tolerance for modifications of the Tyr ring (teicoplanin-type permissive pockets), tolerance for β-hydroxylation (balhimycin-type permissive pockets) and those with stringent specificity for Tyr (specific pockets). Important changes in the amino acids within the specificity code are coloured, with those conserved with Tyr-specific pockets shown in green, changes seen in Tyr-permissive pockets coloured yellow (teicoplanin-type) and grey (balhimycin type) and the rufomycin A3 domain shown in purple; of note in this regard in the Cys to Gly alteration at position 9 of the specificity code.

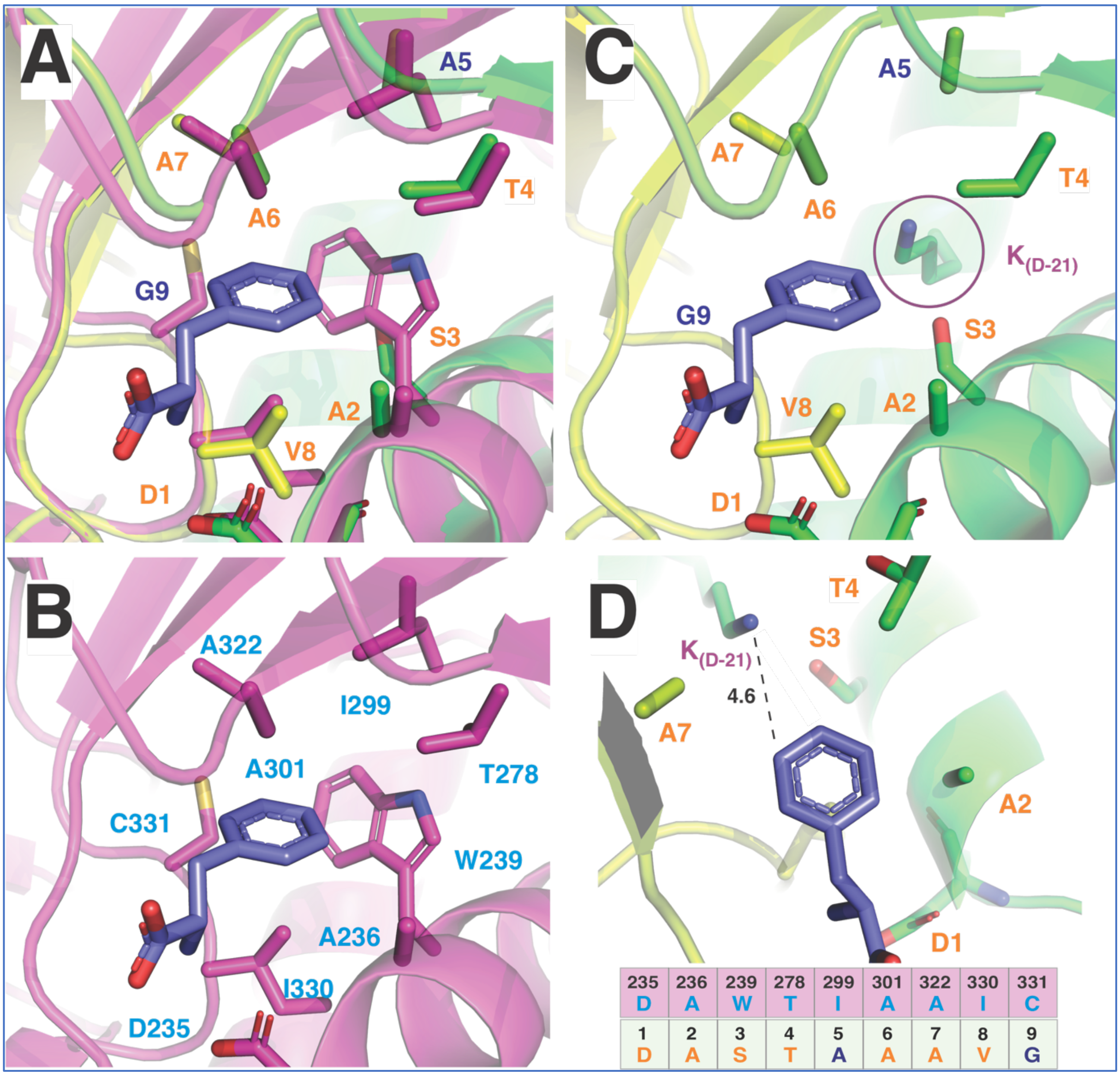

**Supplementary Figure S22.** Structural analysis of the RufT A3 selection pocket using an AlphaFold mode (*77*) in comparison with the phenylalanine activating A-domain Phe (PDB: 1AMU, Chain A). (*78*) Overlay of the structure (A) with the selectivity encoding residues indicated for RufT (A). Structure of PheA in complex with Phe, with selectivity encoding residues numbered (B). Predicted structure of the RufT A3 pocket from an AlphaFold model indicating the key residues in the specificity code together with the unusual Lys residue found at D-21 (C). An alternate view of the RufT A3 domain model showing the distance (in Angstrom) from $Lys_{D-21}$ to the *m*-position of Phe in the PheA structure (D). Specificity code residues are list at the bottom right. PheA structure shown in magenta, with phenylalanine shown in blue sticks; RufT A3 model is shown as Jones' rainbow from N- to the C-termini.

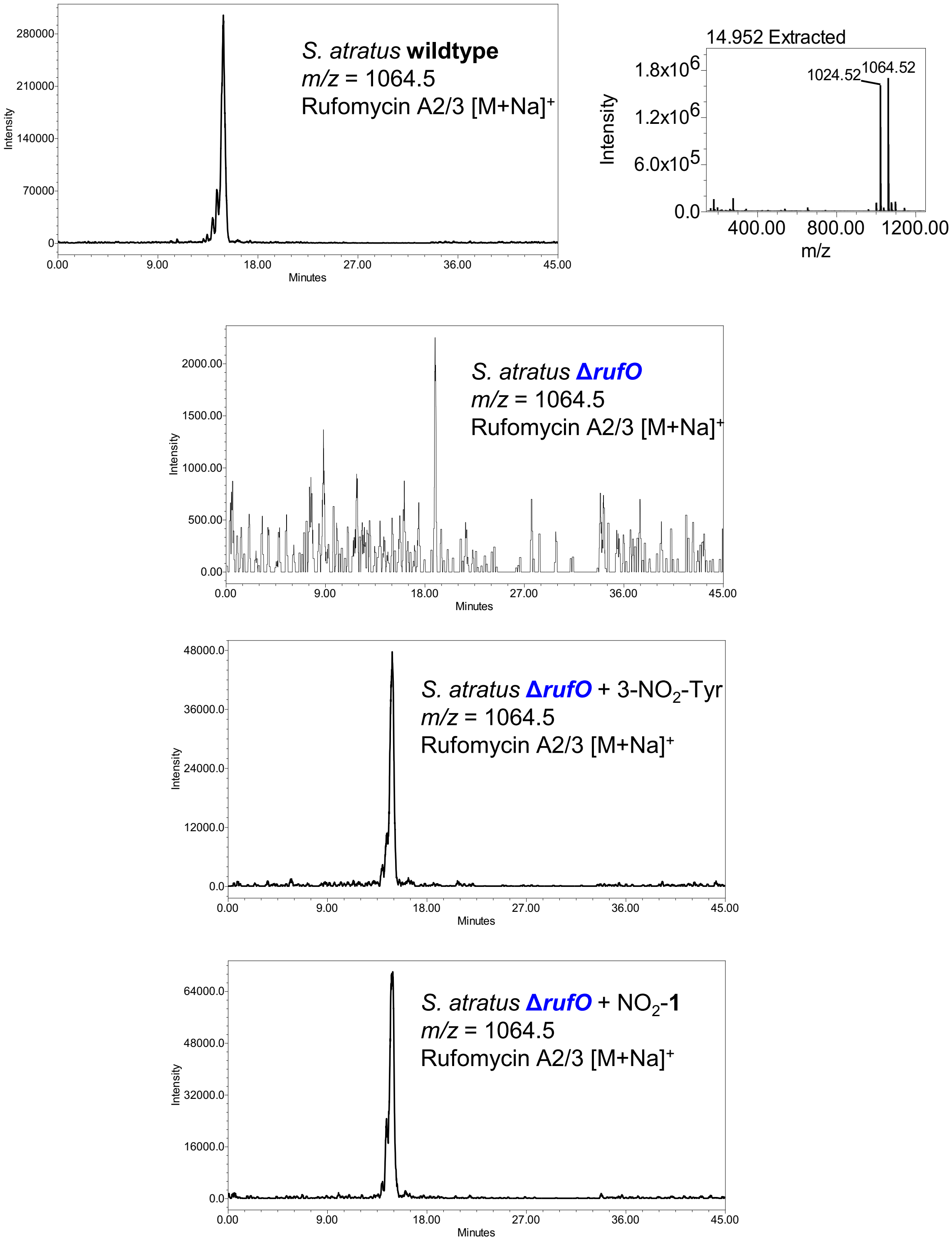

**Supplementary Figure S23. Extracted Ion Chromatogram (EIC) data testing rufomycin production in *S. atratus* wildtype and Δ*rufO* mutants with and without feeding of 3-$NO_2$-Tyr and $NO_2$-1.** Measured by *m/z* = 1064.5 corresponding to rufomycin A2/3 $[M+Na]^+$. Mass spectrum (top right) displaying the most abundant ions at 14.936 minutes.

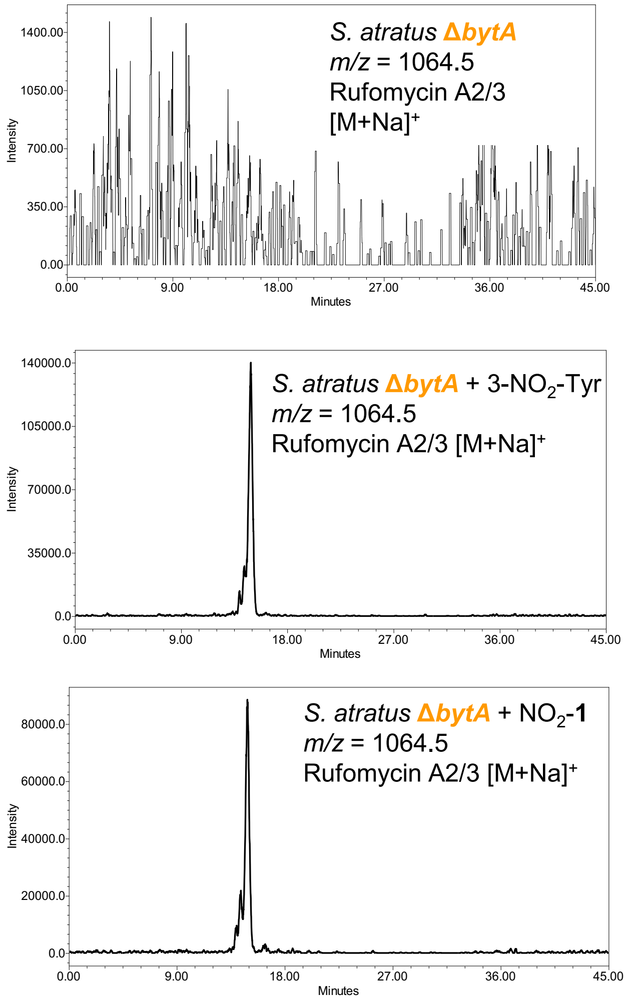

**Supplementary Figure S24. Extracted Ion Chromatogram (EIC) data testing rufomycin production in *S. atratus* Δ*bytA* mutants with and without feeding of 3-$NO_2$-Tyr and $NO_2$-1.** Measured by *m/z* = 1064.5 corresponding to rufomycin A2/3 $[M+Na]^+$.

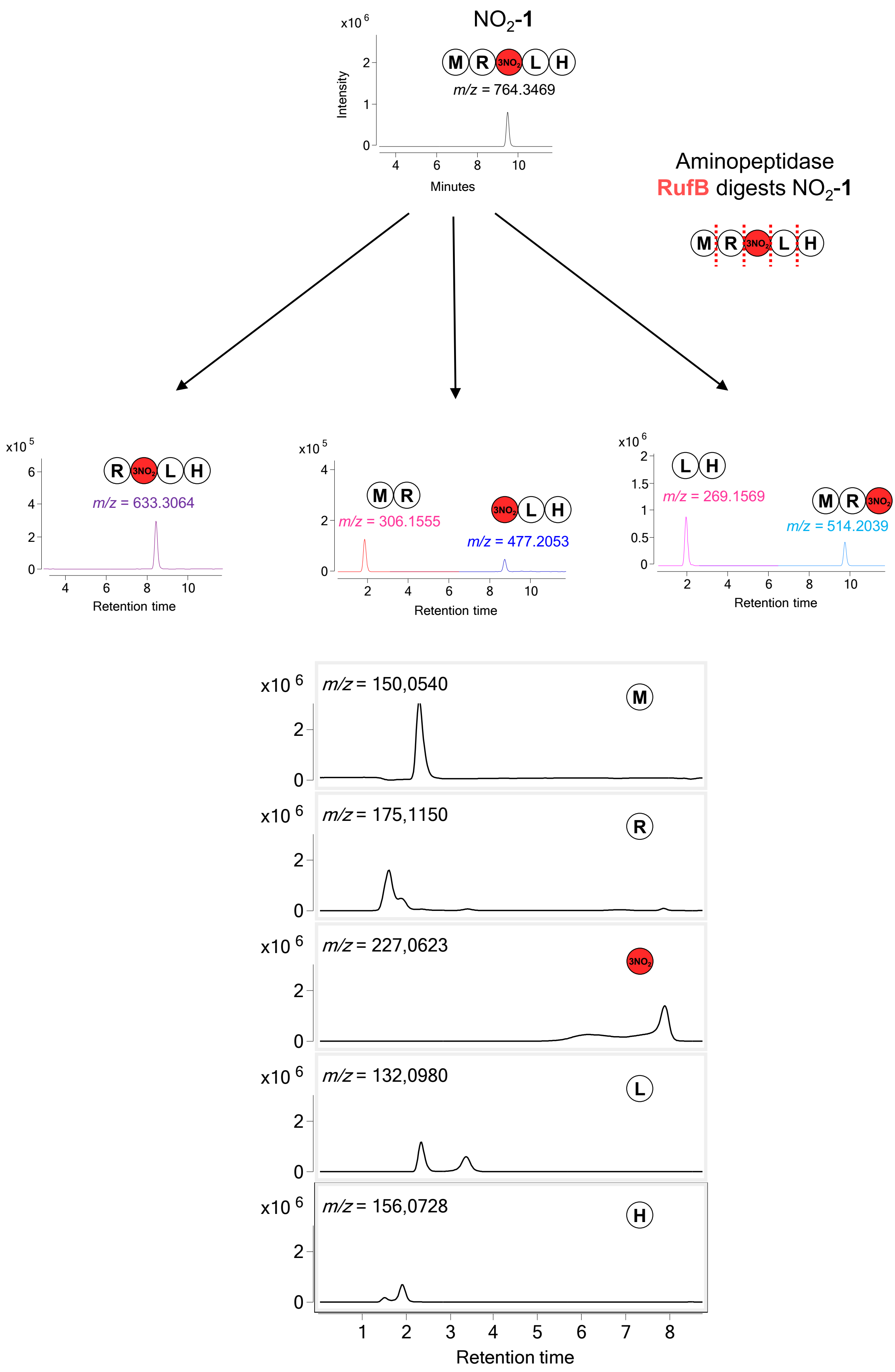

**Supplementary Figure S25**. RufB completely digests the modified pentapeptide $NO_2$-**1**.
